# Who Formed All That Iron?: A Novel Antarctic Chemolithotroph Drives Iron Biomineralization

**DOI:** 10.64898/2026.03.26.714105

**Authors:** Jaekyung Yoon, Boyoung Lee, Kyu-Cheul Yoo, Min-Jung Kwak, Hae Jung Song, Chung Yeon Hwang, Yusook Chung, Kitae Kim, Soon-Kyeong Kwon, Ju Yeon Song, Hwan Su Yoon, Jihyun F. Kim

## Abstract

Iron mineralization has profoundly influenced Earth’s biogeochemical history^1,2^, yet the specific mechanisms underlying banded iron formation (BIF) remain unresolved^3–5^. Here we profile the microbiomes of Holocene sediments beneath the Larsen C Ice Shelf (LCIS), Antarctica^6–8^, through stratigraphic analysis of sedimentary ancient DNA combined with metagenomics. Distinct microbial phases aligned with glacial facies boundaries, with sub-ice shelf communities dominated by chemolithoautotrophs including an uncultured *Thermodesulfovibrionia*. This bacterium, visualized by fluorescence *in situ* hybridization and designated ‘*Candidatus* Mariimomonas ferrooxydans’ (phylum *Nitrospirota*), emerged as a keystone taxon with high network centrality. Its genome encodes Cyc2, a fused porin–cytochrome outer membrane protein implicated in Fe(II) oxidation. Heterologous expression of Cyc2 in *Escherichia coli* confirmed its ability to catalyze iron oxidation, supporting iron precipitation under dark, anoxic conditions. These pristine LCIS sediments, unaltered since the last glacial maximum, provide a modern analogue for synglacial BIFs deposited during Neoproterozoic Snowball Earth events. Our findings deliver direct genomic and functional evidence for chemolithotrophic iron oxidation, challenge phototroph-centric models of BIF genesis, and highlight microbial iron cycling as a recurring force in Earth’s geochemical evolution. Beyond Earth, these insights inform interpretations of iron deposits on other planetary bodies.

## Main

The interplay between inorganic elements and the biosphere has shaped Earth’s crust and atmosphere^9–11^. Iron is central to Earth’s redox balance and biogeochemical cycles, yet the pathways of its oxidation and the microbial agents involved remain unresolved. Banded iron formations (BIFs), with alternating iron- and silica-rich laminae, dominate the Precambrian sedimentary record and constitute a major reservoir of iron ore^1,2,12–14^. Despite decades of study, competing hypotheses persist regarding their genesis, ranging from abiotic oxidation to microbially mediated processes^3–5,15^. Resolving the biological contribution to BIF formation is therefore critical for reconstructing ancient Earth environments and their geochemical evolution.

Although large-scale BIF deposition largely ceased after the Proterozoic, iron mineralization may continue in diverse modern settings, offering analogues for ancient processes. The Holocene, spanning the last ∼11,700 years^15,16^, is often regarded as climatically stable, yet it records substantial sea-level rise and pronounced regional variability^17–20^. Within this epoch, microbial activity under fluctuating redox conditions provides a unique opportunity to examine how iron cycling operates in an oxygenated world. Investigating Holocene sediments can thus illuminate whether mechanisms reminiscent of Precambrian BIF formation persist, bridging the gap between ancient synglacial iron deposition and modern microbial iron mineralization.

As a region of rapid warming^21^, the Antarctic Peninsula serves as a touchstone for climate change and its impact on global ice dynamics. The Larsen C Ice Shelf (LCIS) of the northwestern Weddell Sea has persisted since the Last Glacial Maximum^22^ but has been thinning throughout the Holocene. Pristine since the last glacial period, sediments beneath the LCIS remained unexposed until the recent retreat of the calving front^7,23^. In this study, we examined the stratigraphic sequence of microbiome profiles in a sediment core (GC16B) collected from the Larsen C embayment on the inner continental shelf^6,8^. This core uniquely preserves an uncontaminated Holocene paleoenvironmental record, offering a rare archive to probe anoxic geochemistry and its relevance to ancient iron deposition.

### Microbiota reflect environmental change

History of the LCIS sediment core spans the Holocene from the grounded glacier through floating ice shelf (11,577 ± 479 calibrated years before present at 192 cm) to consolidated sea-ice conditions^6–8,24^. Microbial community profiles of GC16B in 37 sedimentary layers (0‒181 cm) were analysed using amplicon sequencing data of the 16S ribosomal RNA gene (Extended Data Table 1). Throughout the layers, dominant prokaryotic phyla in the LCIS sediment layers included *Chloroflexi*, *Proteobacteria*, *Planctomycetes*, *Acidobacteria*, ‘*Candidatus* Patescibacteria’, ‘*Candidatus* Atribacteria’, and *Nitrospirae* (Fig. 1a). *Euryarchaeota* was dominant in three layers collected from 170‒181 cm. When examined at the family level, succession over three stages was obvious. For example, ‘*Candidatus* Methanoperedenaceae’ (average relative abundance 47.9 ± 12.4%) was dominant in the layers of 170‒181 cm but diminished above 155 cm. Other major families in these layers included *Burkholderiaceae* (8.4 ± 2.7%), an uncultured family in *Thermodesulfovibrionia* (3.6 ± 1.7%), an unassigned family in *Deltaproteobacteria* (3.2 ± 1.6%), an unassigned family in *Chloroflexi* (3.1 ± 1.0%), and *Desulfobacteraceae* (3.1 ± 1.0%). Dominant families in the layers of 45‒166 and 0‒41 cm are shown in Fig. 1a and Extended Data Fig. 1a, b.

**Fig. 1.**
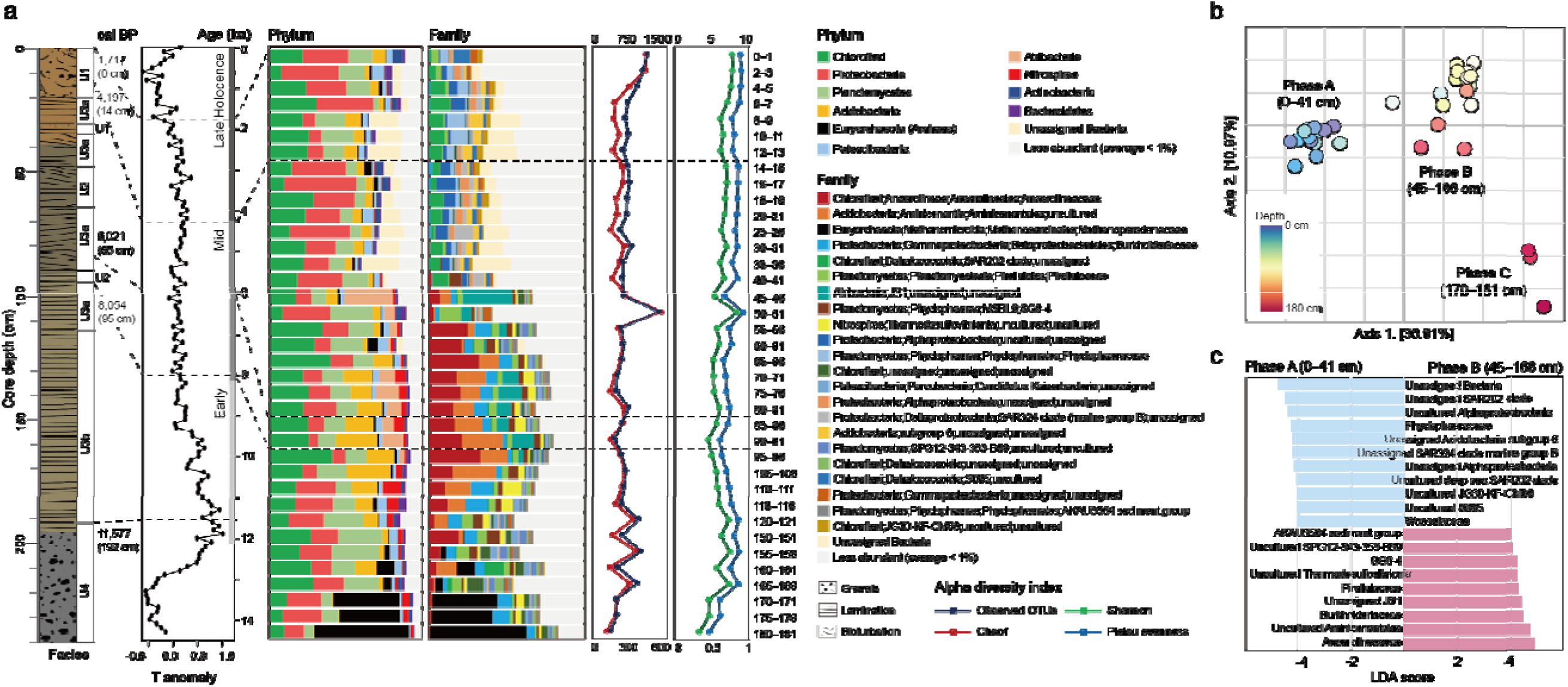
Microbiota structure of the Holocene sediment beneath the Larsen C Ice Shelf. **a,** Sedimentary facies and climate events of GC16B. Four lithostratigraphic intervals are recognized: glacial till diamicton (U4; 192 cm to bottom), a parallel-laminated mud (U3b; 117‒192 cm), cross-laminated mud (U3a; 94‒117, 63‒90, 34‒47, 21‒32 cm), faintly cross-laminated mud (U2; 90‒93, 47‒63 cm), and sandy to homogenous mud with ice-rafted debris (U1; 32‒34, 0‒21 cm). cal BP, calibrated years before present; ka, thousand years. **b,** Principal coordinates analysis plot of the amplicon sequences at the OTU level using the weighted UniFrac distance matrix. Layers cluster into three phases along the depth. Phase A (0‒41 cm) roughly corresponds to oxygenated and bioturbated sandy clay under occasionally open marine conditions; phases B‒C (45‒181 cm) reflect well-laminated iron-containing crystalline silty clays with no diatoms under the floating ice shelf. **c,** Families determined by LEfSe that most likely explain differences between phases A and B. Those having an LDA value of ≥4 are shown in the figure. Most are unassigned or uncultured.

Calculation of sample variance between the layers using the weighted UniFrac distance matrix at the operational taxonomic unit (OTU) level revealed that the community profiles could be grouped into three distinct phases (Fig. 1b). Hierarchical clustering with single-linkage also resulted in three distinct clusters: the uppermost facies from the top to 41 cm designated as phase A (15 samples), the facies spanning 45‒166 cm as phase B (19 samples), and the facies spanning 170‒181 cm as phase C (3 samples). Division of phases B and C was attributed to the dominance of ‘*Ca.* Methanoperedenaceae’ in phase C, although they share *Burkholderiaceae* and an uncultured *Thermodesulfovibrionia* among others. Unconformity of the microbiota profiles between the phases coincides with LCIS retreat. Phase A roughly corresponds to homogenous mud with ice-rafted debris under consolidated pack-ice conditions (occasionally open marine), and phase B and C are laminated iron-containing crystalline facies without diatoms under the floating ice shelf^6–8,24^. These results suggest that the microbiota profiles of the LCIS sediment reflect the marine environment when it was deposited.

We then conducted linear discriminant analysis (LDA) effect size (LEfSe)^25^ evaluation between phases A and B to identify the families that were responsible for the differences between the open or closed marine settings (Fig. 1c). LEfSe identified an unassigned *Bacteria*, an unassigned *Dehalococcoidia* SAR202 clade, an uncultured *Alphaproteobacteria*, *Phycisphaeraceae*, an unassigned *Acidobacteria* subgroup 6, and others as discriminative families with LDA scores ≥4 in phase A. In phase B, *Anaerolineaceae*, an uncultured *Aminicenantales*, *Burkholderiaceae*, an unassigned ‘*Ca*. Atribacteria’ JS1, *Pirellulaceae*, an uncultured *Thermodesulfovibrionia*, and *Phycisphaerae* SG8-4 were among the discriminative taxa. These results further confirmed the microbiota profiles based on relative abundance as shown in Fig. 1a, and the abundance distribution of dominant families (Extended Data Fig. 1c). Alpha diversity was stable in phase A with slight increase toward the top layers and some fluctuations were apparent in phase B (Fig. 1a). Compared to phase B, phase A exhibited significantly higher observed OTUs (*P* = 0.011), Pielou’s evenness (*P* = 0.00037), and Shannon (*P* = 0.00059).

### Co-occurrence pinpoints keystone taxa

To dissect the potential relationships among microbial members in the LCIS sediment communities, their co-occurrence was explored. Construction of the correlation network at the family level revealed the division of the communities into two sub-networks (Fig. 2a). One sub-network, composed of 55 families abundant in phase A, was in the open marine setting; the other, comprising 52 families abundant in phases B and C, was covered by the ice shelf. ‘*Ca.* Methanoperedenaceae’ formed a small sub-hub in the phase B‒C sub-network. This result suggests that microbial members thriving in different marine environments rarely co-exist.

**Fig. 2.**
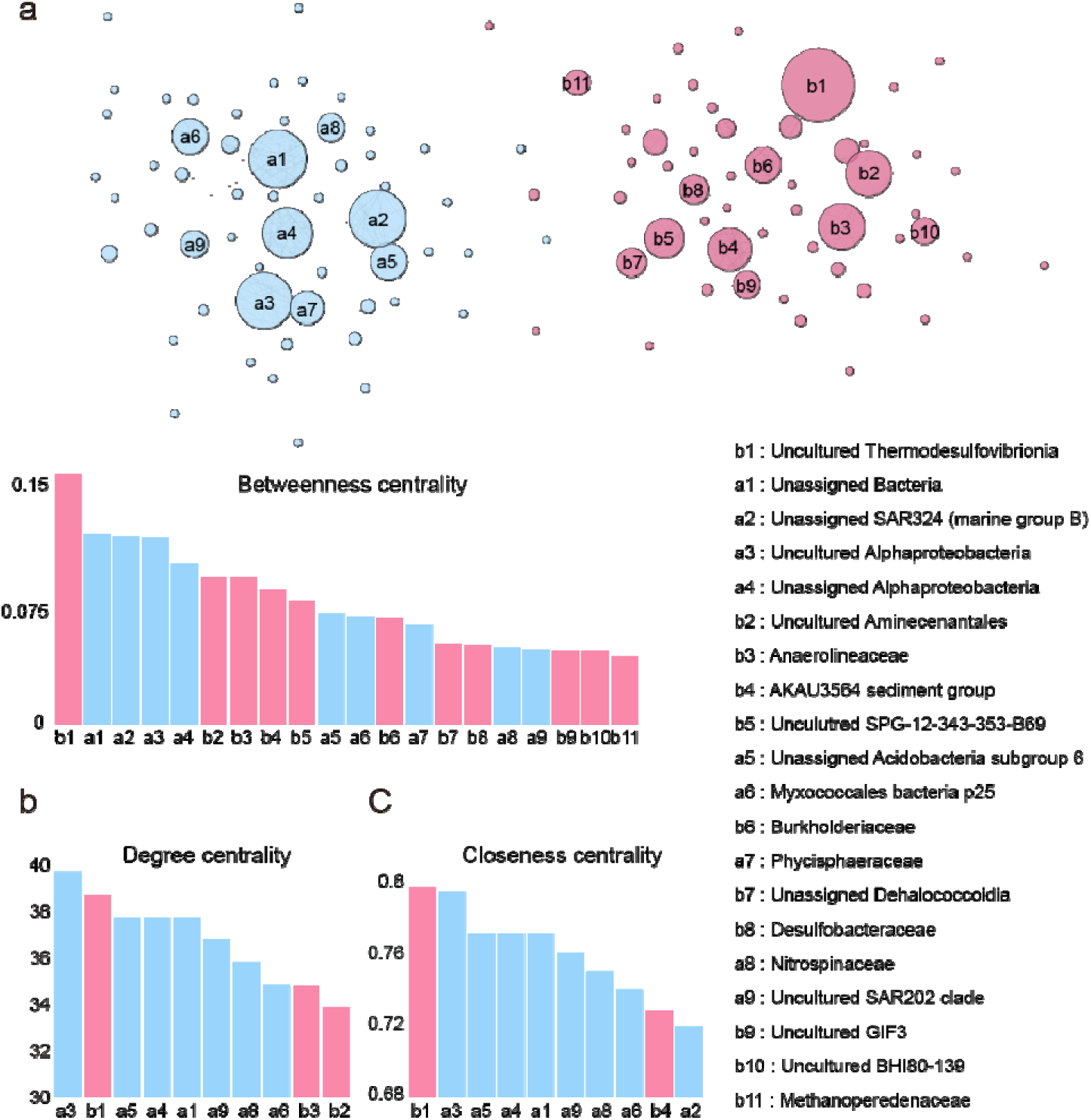
Co-occurrence network of the Larsen C Holocene sediment microbiota. Correlation network of the microbial communities at the family level was constructed using SparCC with the correlation coefficient cut-off value of 0.5 to infer ecological associations between microbial members based on taxonomic composition. Centrality: **a,** betweenness; **b,** degree; **c,** closeness. An uncultured *Thermodesulfovibrionia* exhibits the highest betweenness, degree, and closeness centrality values in the phase B‒C sub-network.

Families exhibiting high network centrality statistics are expected to play important roles in the community. Those with high centrality values of betweenness, degree, and closeness included an unassigned *Bacteria,* an uncultured *Alphaproteobacteria*, and an unassigned *Alphaproteobacteria* in the phase A sub-network, and an uncultured *Thermodesulfovibrionia*, *Anaerolineaceae*, and an uncultured *Aminicenantales* in phase B‒C (Fig. 2a–c). Families with high centrality statistics had relatively similar values in the phase A sub-network, suggesting that they share key roles in the community. In phase B‒C, however, an uncultured *Thermodesulfovibionia* exhibited markedly higher centrality values, indicating that it drives the microbiome structure and possibly function, whereas others have supportive roles.

Further analysis of the phase B–C sub-network revealed three distinct clusters. Cluster 1 was composed of an uncultured *Thermodesulfovibrionia* and the AKAU3654 sediment group, whereas Cluster 2 included *Anaerolineaceae* and an uncultured *Aminicenantales*. Cluster 3 consisted mainly of *Burkholderiaceae*, an unassigned *Dehalococcoidia*, and *Desulfobacteraceae* (Extended Data Fig. 2a). Clusters 2 and 3 exhibited a strong negative correlation (Spearman’s ρ = −0.76, p < 0.0001), suggesting that these two clusters alternately dominated the community within phase B (Extended Data Fig. 2b). Accordingly, the interaction patterns between Cluster 1 and the other clusters may have varied depending on the relative dominance of Clusters 2 and 3. Cluster 2, which included fermentative taxa such as *Anaerolineaceae*, was characterized by taxa commonly associated with the degradation of complex organic matter and the production of metabolic intermediates. In contrast, Cluster 3 was characterized by the presence of sulfate-reducing bacteria, including *Desulfobacteraceae*, which are frequently observed under strongly reducing conditions in marine sediments. Notably, previous study has reported associations between an unassigned *Dehalococcoidia*, *Desulfobacteraceae,* and iron-reducing processes^6^, suggesting that intervals dominated by Cluster 3 may reflect sedimentary conditions compatible with enhanced iron reduction.

Functional profiles of the microbial communities were predicted using PICRUSt2^26^, despite uncertainties related to a number of unclassified or uncultivated taxa. The profiles at the pathway levels were distinctive between the three sediment phases, with 45 pathways significantly differing (FDR < 0.0005) between the two marine environments; i.e. oxygenated open marine or anoxic sub-ice shelf (Extended Data Fig. 3). The succession of metabolic profiles was apparent with abrupt changes at the boundaries: 34 pathways were enriched in phase A and 11 in phases B‒C. Metabolic pathways dominant in phase A encompassed six metabolism categories; the inclusion of active carbon fixation and oxidative phosphorylation suggest that biomass production prevailed. Increased δ^13^C, δ^15^N, and biogenic opal were observed for phase A^6,8^. In contrast, a limited number of metabolic pathways including biosynthesis of sphingolipids, glycans, and flavonoids, and degradation of aromatic hydrocarbons were more prevalent in phases B‒C. Methane metabolism being specifically enriched in phase C is likely due to ‘*Ca.* Methanoperedenaceae’.

### Metagenome reveals anoxic ecophysiology

To uncover the gene repertoire of the LCIS microorganisms, we further performed whole genome amplification followed by whole-metagenome shotgun sequencing using the Ion Torrent PGM and Illumina HiSeq platforms for selected layers spanning the sediment top to 236 cm (Extended Data Table 1). Metagenomic reads were then assembled and binned to reconstruct metagenome-assembled genomes (MAGs)^27^. For the former datasets, co-assembly of all the reads yielded a medium-quality (≥50% completeness, ≤10% contamination)^28^ genomic bin (LCMS_PCM_Al; Extended Data Fig. 4) that matches *Anaerolineaceae* in the 16S rDNA data. Additionally, 20 medium-quality MAGs were binned from the HiSeq contigs.

General metabolic functions of the MAGs were then deduced; Extended Data Fig. 4 shows the percentage of genes present for pathway modules in energy metabolism that are important for biogeochemical cycles in marine environments. In comparison to those in phase A, MAGs enriched in phase B contained numerous genes in the ancient Wood–Ljungdahl pathway, a pivotal carbon fixation pathway under anoxic conditions^29^. Moreover, seven MAGs included a gene encoding carbon monoxide dehydrogenase, a key enzyme in the pathway. Thus, bacteria with these MAGs may play pivotal roles in the carbon cycle to supply feedstock to the community. With regard to methane metabolism, the MAGs contained high percentages of genes in formaldehyde assimilation, except for the serine pathway. Anaerobic energy metabolism, including the Wood–Ljungdahl pathway, nitrogen fixation, and dissimilatory sulphate reduction, appeared active based on phase B-assembled contigs (Extended Data Table 2).

### Iron oxidation by a keystone bacterial taxon

Notably, LCMS-H23C-Mi, LCMS-H26C-Mi, and LCMS-H39C-Mi, which matched an uncultured *Thermodesulfovibrionia*, each encoded a putative Fe(II) oxidase, Cyc2 (Fig. 3a). Cyc2 constitutes a fused porin-cytochrome in the outer membrane that acts as the primary electron acceptor for the respiratory chain^30,31^. LCMS-H39C-Mi contained the full-length Cluster 3-type Cyc2 including the signal peptide, the cytochrome *c* domain with a CXXCH heme-binding motif, and the transmembrane β-barrel domain with 16 antiparallel β-strands (Extended Data Fig. 5a–c). N-terminal sequences were found in LCMS-H23C-Mi and LCMS-H26C-Mi. These three were most similar to a hypothetical protein in *Deltaproteobacteria* SURF_14. Phase B-assembled contigs also contained parts of a Cluster 2-type Cyc2 orthologous to a *Nitrospirae* UBA9159 protein. A neutrophilic Fe(II) oxidation pathway involving Cyc2 has been recently proposed for microaerophilic *Zetaproteobacteria*, in which electrons may pass through the diheme cytochrome Cyc1 or other periplasmic electron carriers to terminal oxidases^31^. Homologs encoding several membrane-bound or periplasmic *c*-type cytochromes and several [Fe-S] proteins that could play roles in electron transfer were found in the three MAGs (Extended Data Table 3). Moreover, homologs encoding dissimilatory or periplasmic nitrate reductase complexes were present in these MAGs, consistent with the reported expression in *Gallionellaceae* sp. of extracellular electron transfer (Cyc2/MtoAB/MofABC) pathways, along with other cytochromes and nitrate reductases, capable of supporting autotrophic Fe(II) oxidation coupled to nitrate reduction in anoxic neutrophilic enrichment culture KS^32^.

**Fig. 3.**
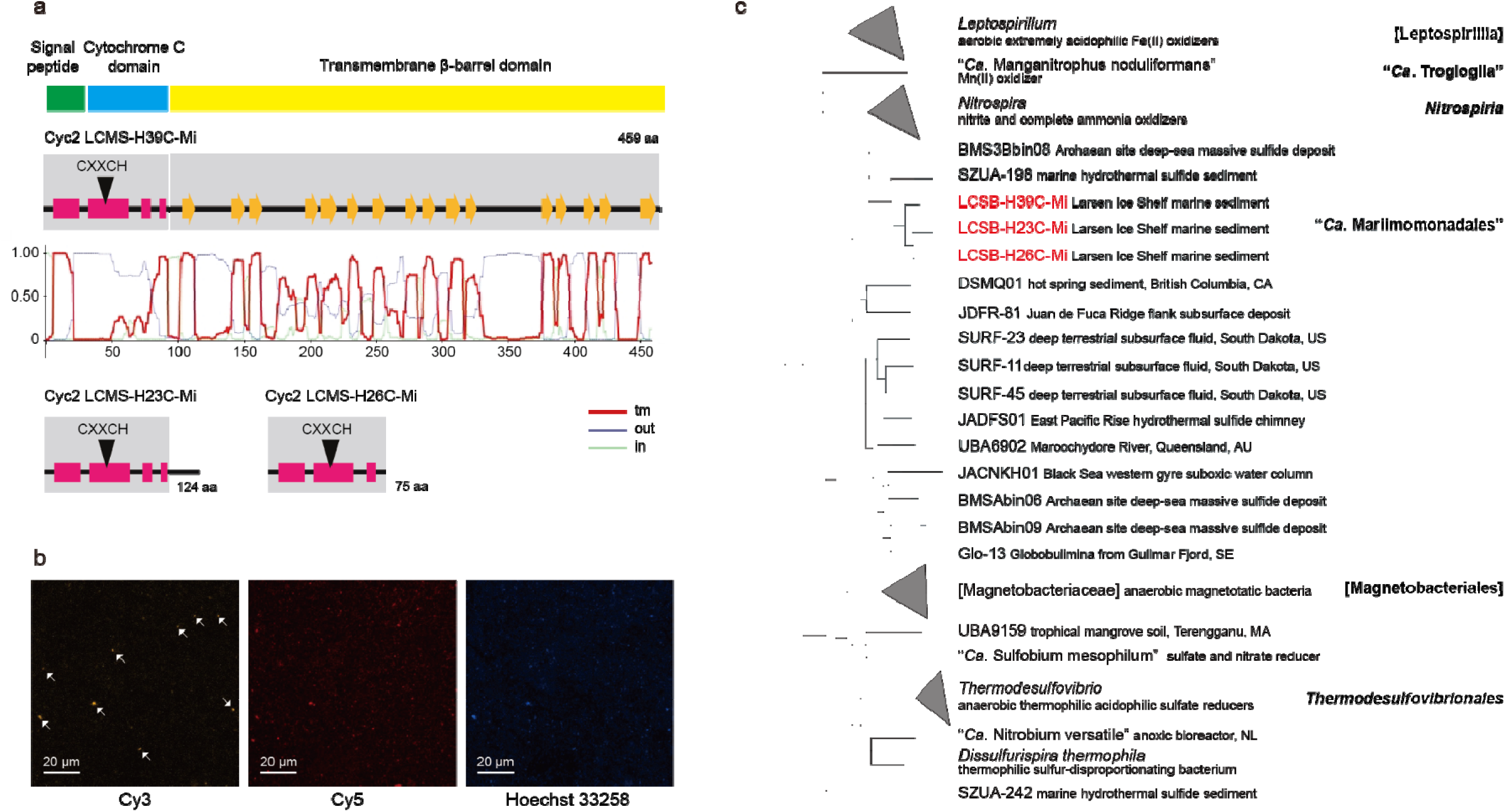
Metagenomics-guided identification of a keystone bacterium, ‘*Candidatus* Mariimomonas ferrooxydans’. Whole metagenomes in select layers spanning the sediment top to 236 cm were subjected to amplification and shotgun sequencing, from which metagenomic reads were assembled into contigs and binned to reconstruct metagenome-assembled genomes. **a,** A putative Fe(II) oxidase Cyc2 identified from three MAGs that match an uncultured *Thermodesulfovibrionia*. LCMS-H39C-Mi from layer 160‒161 cm encodes full-length Cyc2 consisting of the signal peptide, the cytochrome *c* domain with a CXXCH haeme-binding motif, and the outer-membrane porin domain with 16 β-strands. N-terminal sequences were found in LCMS-H23C-Mi (80‒81 cm) and LCMS-H26C-Mi (95‒96 cm). tm, transmembrane; out, extracellular; in, periplasmic region. **b,** Fluorescence *in situ* hybridization images of the LCIS sediment sample from 70‒71 cm. Oligonucleotide probes target ‘*Ca*. M. ferrooxydans’ (Mari_866F_Cy3, orange) and general bacteria (EUB338_Cy5, red). DNA was stained with Hoechst 33258 (blue). Scale bar, 20 μm. **c,** Phylogeny of the phylum *Nitrospirae* (*Nitrospirota* 2018), which contains *Nitrospira*, *Leptospirillum*, *Thermodesulfovibrio*, magnetobacteria, and a novel order ‘*Ca*. Mariimomonadales’. Using *Desulfovibrio vulgaris* as an outgroup, 400 protein markers from 46 *Nitrospirota* isolate genomes and MAGs were selected and aligned. Posterior probabilities of all branches were 1.0.

Similar to the output from microbiota analysis for an uncultured *Thermodesulfovibrionia* exhibiting <87% 16S rDNA identity with any cultivated members of the phylum *Nitrospirota*, which contains *Nitrospira*, *Leptospirillum*, *Thermodesulfovibrio*, and ‘*Candidatus* Magnetobacterium’, GTDB^33^ assigned LCMS-H23C-Mi, LCMS-H26C-Mi, and LCMS-H39C-Mi (97.10–97.51% nucleotide identities; Extended Data Fig. 6) to an unclassified order in the class *Thermodesulfovibrionia*. The associated microbes appear to represent a chemolithoautotroph based on their ability to fix carbon, reduce nitrate and sulphate, and oxidize iron (Extended Data Table 3). Notably, similar to a recent report^34^, in which a *Thermodesulfovibrionia* bacterium is a key player in nitrogen and sulphur cycling, it occupies an important place in the co-occurrence sub-network of phase B–C with markedly high centralities (Fig. 2). Fluorescence *in situ* hybridization-mediated visualization (Fig. 3b and Extended Data Fig. 7) of 16S rRNA-targeted and fluorophore-labelled cells confirmed the presence of an uncultured *Thermodesulfovibrionia*. We propose the epithet ‘*Candidatus* Mariimomonas ferrooxydans’ for these yet-uncultured bacterial strains (taxonomic proposal presented in Supplementary Information) based on their genomic characteristics. No reference genomes are closely related to this provisional species; moreover, its genome shared just 72% average nucleotide identity to BMS3Bbin08 (*Nitrospirae* Clade B) of an as-yet-uncultivated chemosynthetic colonizer in seafloor massive sulphide deposits^35^. Several related MAGs have also been recovered from subsurface fluids and other sources. Together, these formed a novel *Candidatus* order ‘Mariimomonadales’ in *Nitrospirota*, distinct from *Thermodesulfovibrionales* or [Magnetobacteriales] (Fig. 3c). These MAGs encode the Wood–Ljungdahl pathway, a nitrate reductase, and the Dsr-Apr-Sat system (Extended Data Table 3), suggesting that the *Candidatus* order overall also maintains a chemolithoautotrophic lifestyle.

### Cyc2 expressed in *E. coli* oxidizes Fe(II)

To confirm the Fe(II) oxidase activity of ‘*Ca*. M. ferrooxydans’ Cyc2 through heterologous expression in *Escherichia coli* ^36^, we chemically synthesized the coding sequence with several modifications (Extended Data Fig. 8): codon optimization, the N-terminal signal peptide replaced with that of OmpA, and addition of the C-terminal Strep-tag II. After cloning of the modified sequence, designated *cyc2*\*, its expression was successfully induced in *E. coli* BL21(DE3)(pEC86)(pET21a-*cyc2*\*) (Fig. 4A). As expected, the Cyc2*-expressing culture of *E. coli* indeed exhibited significantly accelerated Fe(II) oxidation compared to that of the control (Fig. 4b), highlighting the identity of ‘*Ca*. M. ferrooxydans’ as an iron-oxidizing bacterium and the feasibility of iron precipitation by Cyc2-catalysed oxidation followed by diagenesis to BIFs. It is worth noting that increased illite crystallinity and Fe/Ti under the anoxic condition have been observed in this sediment^6,8^, which is consistent with the abundance distribution of an uncultured *Thermodesulfovibrionia* (Extended Data Fig. 1c).

**Fig. 4.**
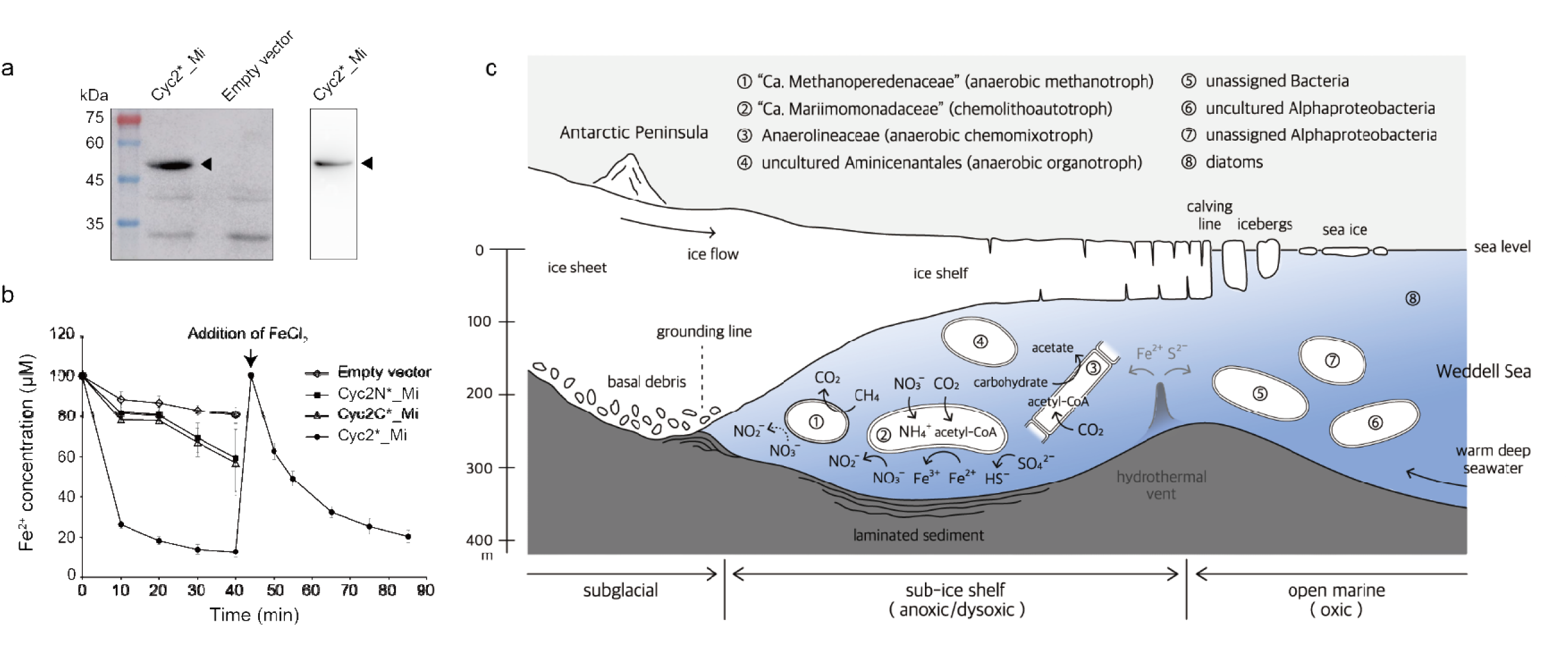
Biomineralization by iron oxidation under the Antarctic ice shelf. **a,** Heterologous expression of ‘*Ca*. M. ferrooxydans’ *cyc2* in *E. coli*. Western blotting (left panel) shows *cyc2\** expression at ca. 49 kDa (arrow head). Haeme staining (right) indicates successful incorporation of haeme to Cyc2*. **b,** Fe(II) oxidation activity of *E. coli* expressing the *cyc2* gene of ‘*Ca*. M. ferrooxydans’. Cyc2*_Mi, *E. coli* BL21(DE3)(pEC86)(pET21a-*cyc2*\*) (full-length Cyc2); Cyc2N_Mi, *E. coli* BL21(DE3)(pEC86)(pET21a-*cyc2N*\*) (N-terminal cytochrome *c* domain of Cyc2); Cyc2C_Mi, *E. coli* BL21(DE3)(pEC86)(pET21a-*cyc2C*\*) (C-terminal outer-membrane porin domain of Cyc2); Empty vector, *E. coli* BL21(DE3)(pEC86)(pET21a) (empty vector). Error bar indicates standard deviation of three biological replicates. **c,** A simplified view of the marine microbial ecosystems recorded in the LCIS Holocene sediment. Keystone taxa in phase A include an unassigned *Bacteria,* an uncultured *Alphaproteobacteria*, and an unassigned *Alphaproteobacteria*; those in phase B‒C include ‘*Candidatus* Mariimomonadaceae’, *Anaerolineaceae* (filamentous), and an uncultured *Aminicenantales* (OP8). ‘*Ca*. Methanoperedenaceae’ (ANME-2d) forms a small sub-hub in the latter. MAG information (‘*Ca*. Methanoperedenaceae’, LCMS-H39C-Mp; ‘*Ca*. Mariimomonadaceae’, LCMS-H39C-Mi; *Anaerolineaceae*, LCMS-H26M-Al), as well as established knowledge on the taxa, was used to infer metabolism. Dotted arrow indicates an unconfirmed pathway. Microbes are exaggerated in size. Illustration by Irin M. Kim, Seoul Women’s University.

## Discussion

Our findings highlight how LCIS sediment preserves a dynamic record of microbial community shifts tightly linked to environmental changes throughout the Holocene. Metagenomic analysis revealed several dominant taxa clustered into three distinct phases that corresponded to geological facies boundaries. The open marine setting (phase A) harbored higher taxonomic richness, whereas the sub-ice shelf sediments (phases B and C) were primarily anoxic and supported diverse chemolithoautotrophic metabolisms, including Wood–Ljungdahl, nitrogen fixation, sulphate reduction, and methane cycling. Co-occurrence network analysis further underscored the importance of key families that shape the overall community structure under anoxic conditions, particularly an uncultured *Thermodesulfovibrionia* with potential for iron oxidation. Overall, these results demonstrate that changing oceanographic and glacial regimes across the Holocene influenced the taxonomic and metabolic landscape of the microbial community, which left evidence in the alterations of the sedimentary layers.

The findings from sub-ice shelf sediments prompt us to re-examine a longstanding geological puzzle: the origin of BIFs. BIFs represent one of the great enigmas in Earth history. Despite decades of intensive investigation, the development of plausible models for the deposition pathway of BIFs has proven challenging because of the rarity of modern ferruginous analogues^37^ and the paucity of biosignatures including microfossils, which are often inconclusive^38^, in (thought to be microbially induced) sedimentary structures. In this work, we addressed the issue through stratigraphic microbiome profiling of the sedimentary ancient DNA deposited beneath the Antarctic ice shelf during the Holocene and by functional validation of the identified Fe(II) oxidase, Cyc2, which provides insights into the ecophysiology and biogeochemistry concerning iron formations.

Popular BIF genesis models posit cyanobacteria and/or anoxygenic photoferrotrophs as the biogenic mechanisms that underpin iron deposition^1,39^, although alternative postulates including the possibility of anaerobic chemotrophic Fe oxipdation, still unknown in modern systems^3^, were recently proposed^40,41^. Our results pinpoint the occurrence of biomineralization by iron bacteria beneath the floating ice shelf, under oligotrophic conditions devoid of light and oxygen. The *Thermodesulfovibrionia* bacterium, visualized using fluorescence *in situ* hybridization and designated as ‘*Ca*. M. ferrooxydans’, formed a novel clade in the phylum *Nitrospirota* along with several yet-uncultivated subsurface chemolithoautotrophs. Its genome unfolded Cyc2, in addition to other cytochromes, [Fe-S] proteins, and nitrate reductases. During the Greenlandian and Northgrippian, ‘*Ca*. M. ferrooxydans’ and other lithoautotrophs^42^ must have utilized Cyc2 to oxidize dissolved Fe(II), possibly originating from subglacial meltwater^43^ or local hydrothermal vents^44^, to produce proton motive force or reducing power (Fig. 4c). This in turn may have nourished a rich anoxic ecosystem of marine plankton^45^ underneath the Antarctic ice shelf. The consequence of Fe(II) oxidization by Cyc2 is the precipitation of Fe(III) due to its insolubility in water. In particular, the Holocene sediment beneath the LCIS is reminiscent^8^ of the synglacial BIFs in the Neoproterozoic ‘Snowball Earth’, featuring dropstone-free laminated sedimentary facies with massive ice-proximal diamictites during the Sturtian glaciation^46^. Furthermore, microbial nitrate-dependent iron oxidation might have been involved in anoxic iron formation, a form of stromatolites found in some of the Earth’s oldest rock formations^47,48^, during much of the Archaean to the early Proterozoic^49^, and even ferric oxide rusty dust on early Mars^50^.

Taken together, our findings demonstrate that Cyc2-bearing chemolithotrophs are sufficient to drive iron biomineralization under anoxic and ferruginous conditions. This functional validation provides direct biological evidence for microbial involvement in BIF genesis, addressing a long-standing debate on their origin. By linking Holocene Antarctic sediments to Precambrian synglacial iron formations, our study highlights how microbial iron oxidation has repeatedly shaped Earth’s geochemical history. Beyond Earth, these insights may also inform interpretations of iron deposits on other planetary bodies.

## Online content

Any methods, additional references, Nature Portfolio reporting summaries, source data, extended data, supplementary information, acknowledgements, peer review information; details of author contributions and competing interests; and statements of data and code availability are available at $$$.

## Acknowledgments

We thank Dianne K. Newman for providing pEC86, Irin M. Kim for depicting schematic illustrations, Jillian F. Banfield and Peer Bork for advices on the manuscript, Sanghoon Kwon and Dong-Woo Lee for discussions regarding BIF and sulphur metabolism, respectively, Eugene Kim for help with English editing, members of the JFK laboratory including HyeonGwon Lee and the HSY laboratory including Duckhyun Lhee for technical assistance, and KOPRI scientists and Araon crew for contributing to the marine geological expedition of the Larsen Ice Shelf (ARA13 Cruise Expedition) in May 2013. This work was financially supported by grants from the National Research Foundation of Korea (RS-2023-NR077253, RS-2024-00401518, RS-2021-NR056580 to J.F.K. and NRF-2022R1A2B5B03002312 to H.S.Y.), Korea Institute of Marine Science & Technology Promotion (RS-2025-02307311 to J.F.K.), and Korea Polar Research Institute (PAP-PD15010) to H.S.Y., C.Y.H., and K.-C.Y. Publication was supported in part by the Brain Korea 21 program, and J.Y. and K.K. are fellowship awardees of this program.

## Author contributions

J.F.K. and H.S.Y. conceived, organized, and supervised the work. K.-C.Y. was responsible for sediment sampling. C.Y.H. performed 16S rDNA sequencing, H.J.S. performed metagenome sequencing, J.Y., M.-J.K., Y.C., and J.Y.S. performed sequence analyses, B.L. conducted fluorescence *in situ* hybridization, and J.Y., B.L., K.K., and S.-K.K. conducted iron oxidation experiments. J.F.K., J.Y., and M.-J.K. interpreted the results. J.F.K. wrote the main text. J.Y., B.L., and H.J.S. prepared Methods, figures, and tables. J.F.K., J.Y., S.-K.K., H.S.Y., and K.-C.Y. edited the manuscript. All of the authors read and approved the final version of the manuscript before submission.

## Methods

### Sample collection

The sediment core was collected from AP LCIS, the north-western part of the Weddell Sea. A whole round core sample of the marine sediment (GC16B; 236 cm below the seafloor at 324 m below the sea level in the approximately 8.7-km-wide basin), mostly consisting of laminated facies, was collected on the northern part of the LCIS embayment (66° 3.89832′ S, 60° 27.69212′ W)^6,8^ (Fig. 1). Sediment ages were estimated by acid insoluble organic and ramped PyrOx 14C dating, and atmospheric temperature anomalies were obtained from the James Ross Island ice core24. Core GC16B is defined by four distinct lithological units from the top (U1) to the bottom. These sedimentary sequences span the transitions from glacial till diamicton under the grounded glacier (U4, 192 cm to bottom) through parallel-laminated mud (U3b, 117‒192 cm; 11,577 ± 479 cal BP at 192 cm)^24^ to cross-laminated mud (U3a, 94‒117, 63‒90, 34‒47, 21‒32 cm; 6,021 ± 429 and 8,054 ± 420 cal BP at 85 and 95 cm, respectively) and indistinct to faintly cross-laminated mud under the floating ice shelf (U2, 90‒93, 47‒63 cm) to sandy to homogenous mud with ice-rafted debris under the consolidated sea ice (U1, 32‒34, 0‒21 cm; 1,717 ± 90 and 4,197 ± 83 cal BP at 0 and 14 cm, respectively). The dark grey sandy diamicton present in U4 was interpreted as rapidly deposited soil proximal to the grounding zone of the ice shelf during the Last Glacial Maximum.

### Microbiome sequencing

Environmental DNA was extracted from 40 layers of the sediment core using the FastDNA Spin Kit for soil (MP Biomedicals, USA). The quantity of genomic DNAs was measured using Picogreen fluorometry. From the top to 20 cm, DNA was extracted at 2 cm intervals and at 5 cm intervals from below 20 cm. A total of 37 layers from the top to 181 cm were selected for amplicon sequencing. The V5–V8 region of the 16S rRNA gene was amplified using universal primers (Uni787F and Uni1391R) and sequenced using pyrosequencing, which produced 2,846± 1,267 reads.

We selected 16 layers distributed evenly throughout the entire core (spanning top to 236 cm; Extended Data Table 1) for whole-metagenome shotgun sequencing. Owing to low DNA concentration (<1 ng for each layer), whole genome amplification was performed using the Yikon MALBAC whole genome amplification kit (Yikon Genomics Co., Ltd., China), which provides a relatively low bias rate through DNA fragmentation^51^. The amplified DNA was purified using the LaboPass™ Tissue mini Kit (Cosmo Genetech, Korea); concentration was determined with a Qubit™ 2.0 Fluorometer (Invitrogen, USA).

Total genomic DNA samples from the 16 layers were subject to shotgun sequencing. DNA shearing was performed to obtain 400-bp fragments, and libraries were constructed using the Ion Plus Fragment Library Kit and Ion PGM Template OT2 400 Kit (Life Technologies, USA). The libraries were sequenced using the ION PGM Sequencing 400 Kit on the Ion Torrent PGM (Life Technologies). The Ion 318™ Chip Kit V2 (Life Technologies) was used for template loading. Library construction and sequencing steps followed the manual provided by Life Technologies. An average of 4,535,898 reads was produced.

Additional sequencing was conducted to produce deep coverage datasets with six layers. Libraries were constructed using the TruSeq Nano DNA prep Kit (350-bp Protocol) and SureSelect/TrueSeq Nano DNA Prep Kit (350-bp Protocol: SureSelect Exome Modify). The libraries were sequenced on the Illumina HiSeq 2500 platform (Illumina Inc., USA) with 2×150-bp paired-ends conducted at DNA Link, Inc. (Korea). An average of 55,922,748 reads was produced.

### Bioinformatics analysis of microbiota data

The QIIME 2 pipeline^52^ was used for microbiota analysis of the pyrosequencing data. Reads were clustered into OTUs with 97% similarity using VSEARCH^53^. Taxonomy assignments of representative sequences were conducted using the Naïve Bayes classifier^54^ trained utilizing the full-length 16S rRNA genes in the Silva 132/99% database^55^. Reads that were assigned as Eukarya, mitochondria, or chloroplasts were removed. Alpha diversity and beta diversity indices were calculated following normalization through rarefaction using a QIIME 2 module. Phylogenetic trees for OTUs in phase A and phase B were deduced after removing the sequences that appeared in fewer than two samples. Reference-based sequence alignment was conducted using SINA-1.7.2^56^ with SILVA_138.1_SSURef_opt.arb as a reference and with the ‘--fasta-write-dna’ option. Aligned sequences were imported into QIIME 2 and masked using the ‘qiime alignment mask’ command. Phylogenetic trees were constructed using iqtree^57^ through the ‘qiime phylogeny iqtree’ command. Among 286 models of nucleotide substitution, the GTR+F+R10 model was selected using the ModelFinder algorithm and phylogenetic distance was calculated. Constructed trees were annotated using iTOL v6.3^58^.

LEfSe^25^ was used for identifying differentially abundant families between phases A and B. Default values were used for analysis: 0.05 for the Kruskal–Wallis test, and 2.0 for the threshold of the logarithmic LDA score. For network construction, the q2-SCNIC plugin^59^ module in QIIME 2 was used. We utilized the SparCC^60^ method for calculating the correlation matrix, and the network was generated using a cut-off value of 0.5. The constructed network was visualized and analysed using Cytoscape 3.8.2^61^. PICRUSt2^26^ was used to predict the functional profiles of sediment microbiota based on 16S rDNA sequences. Functional profiles at the KEGG orthologue (KO) ^62^ level were predicted using the command ‘picrust2_pipeline.py’ version 2.3.0-b, and profiles at the KEGG pathway-level were deduced based on normalized KO profiles using the ‘pathway_pipeline.py’ command.

### Analysis of whole metagenome data

Whole-metagenome shotgun sequence reads were analysed using the metaWRAP v1.2 pipeline^63^. Reads in each sample were trimmed using the metaWRAP-Read_qc module and assembled into contigs using MEGAHIT^64^. Prodigal v2.6.3^65^ was used for predicted coding sequences (CDSs) and GhostKOALA^62^ (last updated May 15, 2019) was used for functional annotated with KOs. KEGG pathway and KEGG module profiles based on KOs were reconstructed using Reconstruct tool in KEGG mapper^66^ (last updated July 1, 2021). Additionally, FeGenie^67^ was used for the identification of iron metabolism-related genes and DiSCo^68^ was used for the identification of sulphur metabolism-related genes.

Genomic bins were obtained using CONCOCT^69^, MaxBin2^70^, and metaBAT2^71^, which were implemented in the binning module of the metaWRAP pipeline. checkM v1.0.18^72^ was used to evaluate the quality of the genomic bins. Reassembly of the genomic bins was performed using the ‘reassembled_bins’ module in the metaWRAP pipeline to improve the quality of the genomic bins. Briefly, reads belonging to each genomic bin were recruited using Bowtie2^73^ with either permissive or strict parameters, and recruited reads were assembled using Spades^74^. Additionally, in the case of some genomic bins with high contamination, recruited reads were assembled using MEGAHIT instead of Spades, and then binning by metaBAT2 was performed to separate the contigs from different microbes. The reassembled genomic bins were evaluated using checkM, and the best quality bins satisfying the medium quality criteria among the reassembled bins and the original bins were selected for further analysis. To recover additional genomic bins, co-assembly with HiSeq sequencing reads of six layers and PGM sequencing reads of 16 layers was performed.

For taxonomic assignments of the genomic bins, the ‘classify_wf’ function in GTDB-Tk^75^ was used incorporating GTDB release95 as the reference database version. For functional annotation of the genomic bins, CDSs were predicted using Prodigal and their functional annotations were determined using BlastKOALA^62^ (last updated May 15, 2019) by searching in the ‘genus_prokaryotes’ KEGG Genes database. KEGG modules were reconstructed from the KO profiles results of BlastKOALA using KEGG mapper. Additionally, genes related to iron metabolism were predicted using FeGenie.

### Analyses of ‘*Ca*. M. ferrooxydans’ MAGs

For further analysis of ‘*Ca*. M. ferrooxydans’ MAGs, we used Prokka 1.14.6^76^, EggNOG mapper v2.0.1^77^, and BlastKOALA^62^ for gene prediction and annotation. In Prokka, Prodigal option –c was set to only call full-length genes containing initiator and terminator codons. Because ‘*Ca*. M. ferrooxydans’ MAGs were of medium quality and included a number of contigs, we removed Prodigal command option –c in Prokka to include partial gene calls. Eggnog-mapper results included eggnog orthologues and several databases including KOs. FeGenie was also applied for the identification of iron-related genes. For comparison of the characteristics of the three ‘*Ca.* M. ferrooxydans’ MAGs, anvio-6.2^78^ was used.

### Construction of the phylogenomic tree

PhyloPhlAn 3.0^79^ was used for construction of the phylogenomic tree of the phylum *Nitrospirae* (*Nitrospirota* 2018)^80^ with 400 universal markers selected by PhyloPhlAn 3.0 using the –d phylophlan parameter ‘–diversity medium –accurate’. The marker gene identification to tree construction steps were performed using the following parameters: Diamond, blastx --quiet --threads 15 --outfmt 6 --more-sensitive --id 50 --max-hsps 35 -k 0; Mafft, --quiet --anysymbol --thread 15 –auto; Trimal, –gappyout; and RAxML, -p 1989 -m PROTCATLG. The Bayesian phylogenomic tree was constructed using MrBayes 3.2.7a^81^ with a mixed amino acid model. The posterior probabilities of all branches were 1.0. The constructed tree was annotated using iTOL v6.

### Fluorescence *in situ* hybridization

Sample fixation and immobilization were performed by referring to the procedure described in recent reviews^82,83^. Briefly, for fixation, the sediment samples were resuspended and incubated in 3% paraformaldehyde buffered with PBS (130 mM NaCl, 7 mM Na_2_HPO_4_, 3 mM NaH_2_PO_4_, pH 7.3) at 4 °C overnight. Bacteria were extracted via Nycodenz separation (Kyongshin Scientific Co., Ltd., Korea) and subsequently immobilized on a GTTP membrane filter (Millipore, Germany) using a vacuum filtration system (Millipore). After permeabilization of the immobilized bacteria on the filter using microwave treatment for 2 min, fluorescence *in situ* hybridization was performed using a 5′-end Cy3-labelled oligonucleotide probe targeting ‘*Ca*. M. ferrooxydans’ (Mari_866F; GGCAATAGTCATCCG) and a 5′-end Cy5-labelled oligonuceleotide probe for general bacteria (EUB338^84^); GCTGCCTCCCGTAGGAGT) in hybridization buffer (0.9 M NaCl, 20 mM Tris-HCl [pH 7.2], 10% dextran sulphate, 1% blocking reagent, 0.01% SDS, and 55% formamide (v/v)) at 40 °C for 3 h. Sequential washing of the filter was performed using washing buffer (0.9 M NaCl, 20 mM Tris-HCl [pH 7.2], 5 mM EDTA, 0.01% SDS, and 55% formamide (v/v)) at 42 °C for 20 min, PBSX (PBS containing 0.05% Triton X-100) at room temperature for 15 min, and distilled water after which the filter was dehydrated with ethanol followed by counterstaining with 2.5 μg/mL 4′,6-diamidino-2-phenylindole (DAPI) or Hoechst 33258 for visualization of general DNA. The images were obtained using a confocal laser scanning microscope (ZEISS LSM900, Germany) equipped with appropriate filter sets for Cy3, Cy5, and DAPI/Hoechst fluorescence. All of the images were at a resolution of 1024 × 1024 pixels.

### Analyses of ‘*Ca*. M. ferrooxydans’ Cyc2

The sequences of putative *cyc2* genes in ‘*Ca*. M. ferrooxydans’ MAGs were analysed. The N-terminal signal peptide was predicted using SignalP-5.09^85^, and the haeme-binding motif CXXCH was confirmed by manual search. The secondary structure of Cyc2 was predicted using PSIPRED 4.0^86^ and Phyre v2.0^87^. The protein structure was predicted using ColabFold^88^, in which AlphaFold2^89^ was implemented. The topology of transmembrane beta-strands was predicted using PRED-TMBB^90^. The phylogenetic tree of Cyc2 was constructed using published sequences^91^. A total of 1593 Cyc2 homolog sequences including functionally confirmed sequences was used. Sequences were aligned using ClustalW^92^ and the phylogenetic tree was constructed with RAxML using the PROTGAMMAWAG model.

### Heterologous expression of Cyc2 in *E. coli*

Coding sequence of the *cyc2* gene of ‘*Ca*. M. ferrooxydans’ was chemically synthesized (Bionics, Korea) with following modifications: codon optimization for optimal expression in *E. coli*, replacement of the signal peptide at the N terminus with that of *E. coli* OmpA, and addition of Strep-tag II to the C terminus. Gene fragments encoding either the N-terminal cytochrome *c* or C-terminal porin domain were also synthesized. The modified sequence of full-length *cyc2*, designated *cyc2*\*, was cloned into the expression vector pET21a and the resulting plasmid pET21a-*cyc2*\* was transformed into *E. coli* BL21(DE3)(pEC86). The plasmid pEC86 is a pACYC184 derivative that constitutively expresses the *E. coli* cytochrome *c* maturation genes *ccmA–H* of the *aeg* operon from the *tet* promoter^93^.

For expression of each protein, *E. coli* BL21(DE3)(pEC86) strains harbouring each expression plasmid were grown to mid-log phase in Lysogeny Broth (LB) medium supplemented with 10mM 2-(N-morpholino)ethanesulfonic acid (MES) and 1 mM aminolevulinic acid, and appropriate antibiotics at 37 °C with vigorous shaking. The *E. coli* cultures were induced with 0.5 mM IPTG and incubated for 16 h at 18 °C with shaking at 250 rpm. Cells expressing Cyc2 turned pink, which indicates the successful incorporation of haeme. The peroxidase activity of cytochrome c was evaluated using an in-gel assay, in which electrophoresis was conducted using the cell lysate. Following electrophoresis, the gel was washed with distilled water for 20 min at room temperature. ECL solution was poured on the gel and chemiluminescence was detected using an Amersham Imager 600 (GE Healthcare Life Sciences, USA).

### Fe(II) oxidase activity assay

Prior to the whole-cell Fe(II) oxidation assay, *E. coli* cells expressing full-length Cyc2 or porin without the cytochrome *c* domain were harvested, washed with sterile LB supplemented with 10 mM MES, pH 6.0, and resuspended into the washed with sterile LB supplemented with 10 mM MES, pH 6.0 at OD_600 nm_ = 2. At 5 min after addition of 100 μM FeCl_2_, the supernatant of the cell suspension was subjected to ferrozine measurement. The supernatant was mixed with 0.2 mM ferrozine and 1.36 M acetate buffer, pH 4.0, and incubated for 15 min in the dark. The absorbance was measured at 562 nm using a microplate reader (Spark^®^, Tecan Trading AG, Switzerland).

### Statistical information

Statistical tests were performed using QIIME2 plugins for microbiota data analysis. LEfSe analysis was conducted using Galaxy implementation (https://huttenhower.sph.harvard.edu/galaxy/). For results of iron oxidation assay, Mann-Whitney *U*-test was used for comparison between two experimental groups, and Kruskal-Wallis was used for comparison between more than two groups.

## Data and code availability

The sequences used in this study were deposited in GenBank under BioProject PRJNA792354, from which sequence information of the 16S rRNA gene and MAGs are available. The accession numbers for LCMS-H23C-Mi, LCMS-H26C-Mi, and LCMS-H39C-Mi of ‘*Ca*. M. ferrooxydans’ are SAMN24803906, SAMN24803907, and SAMN24803905, respectively.

## Extended Data Figures

**Extended Data Fig. 1.**
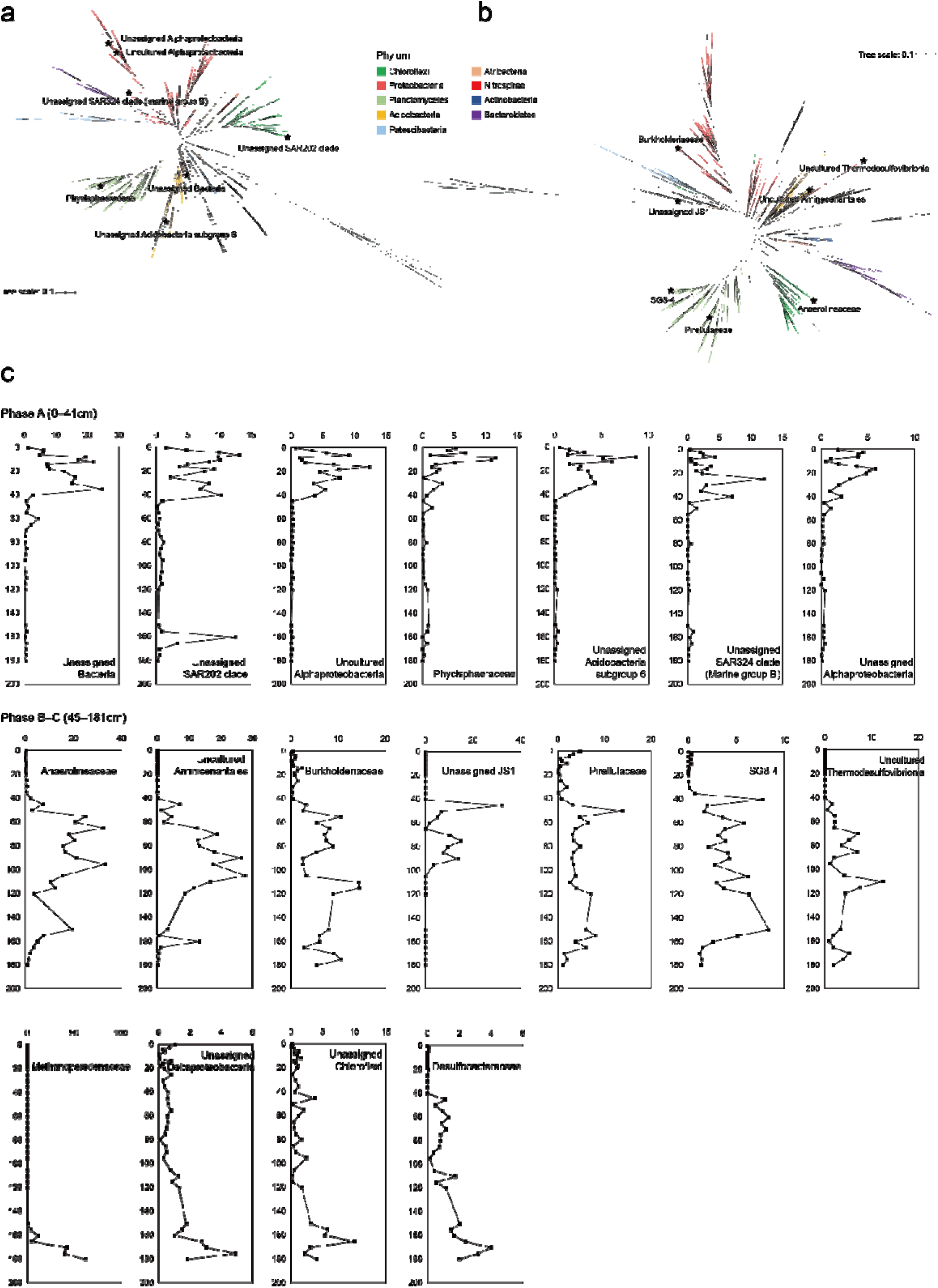
Phylogeny of OTUs and dominant families in the Holocene sediment beneath the Larsen C Ice Shelf. **a,** An unrooted phylogenetic tree of OTUs in layers 0‒41 cm. OTUs appeared less than two samples were removed and the tree was constructed using iqtree on 998 OTUs. Dominant families included an unassigned family in Bacteria (11.2±7.3%), an unassigned family of the SAR202 clade (*Dehalococcoidia*) (7.3±3.3%), an unassigned family in an uncultured order of *Alphaproteobacteria* (5.0±3.2%), *Phycisphaeraceae* (3.9±3.5%), an unassigned family in *Acidobacteria* subgroup 6 (3.3±2.2%), an unassigned family of the SAR324 clade (marine group B; *Deltaproteobacteria*) (3.1±3.0%), and an unassigned family in *Alphaproteobacteria* (3.0±1.7%). Stars indicate representative OTU of those dominant families. **b,** An unrooted phylogenetic tree of OTUs in layers 45‒166 cm. OTUs appeared less than two samples were removed and the tree was constructed using iqtree on 1,459 OTUs. Dominant taxa included *Anaerolineaceae* (15.3±9.2%), an uncultured family in *Aminicenantales* (11.4±8.2%), *Burkholderiaceae* (6.7±3.7%), an unassigned family of JS1 (*Atribacteria*) (5.5±8.2%), *Pirellulaceae* (5.0±2.7%), SG8-4 (*Phycisphaerae*) (3.9±1.8%), and an uncultured family in *Thermodesulfovibrionia* (3.7±3.0%). Stars indicate representative OTU of those dominant families. **c,** Relative abundance distribution of dominant families. Families with the average relative abundance of ≥3% in each phase are shown.

**Extended Data Fig. 2.**
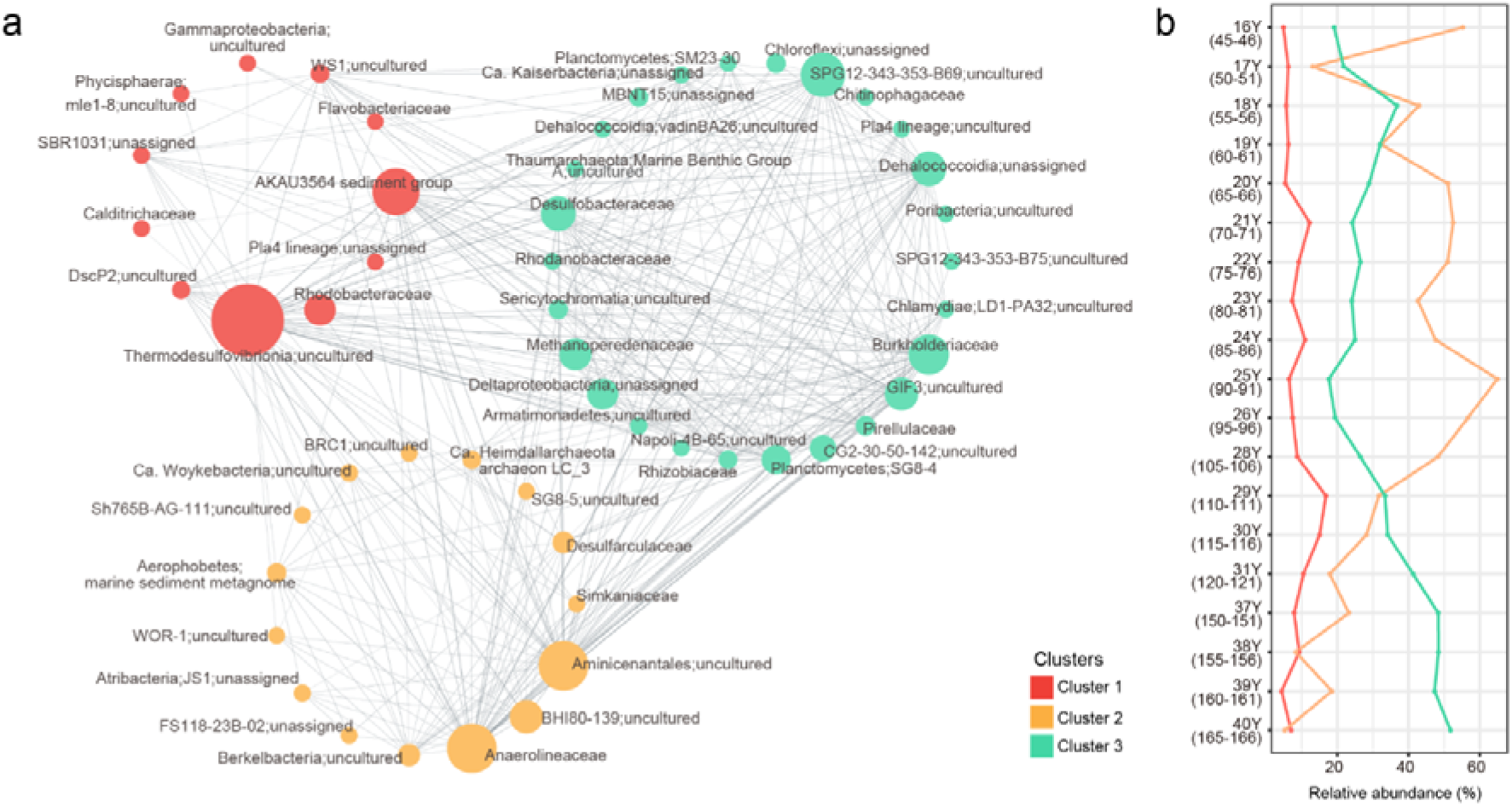
Co-occurrence network representing the sub-network associated with Phase B‒C at the family level. **a,** The sub-network corresponding to Phase B‒C was extracted from the full co-occurrence network. Community structure was determined using the GLay clustering algorithm, which resolved three distinct clusters. Node size reflects each family’s betweenness centrality within the network. **b,** Relative abundances of the three clusters across Phase B samples.

**Extended Data Fig. 3.**
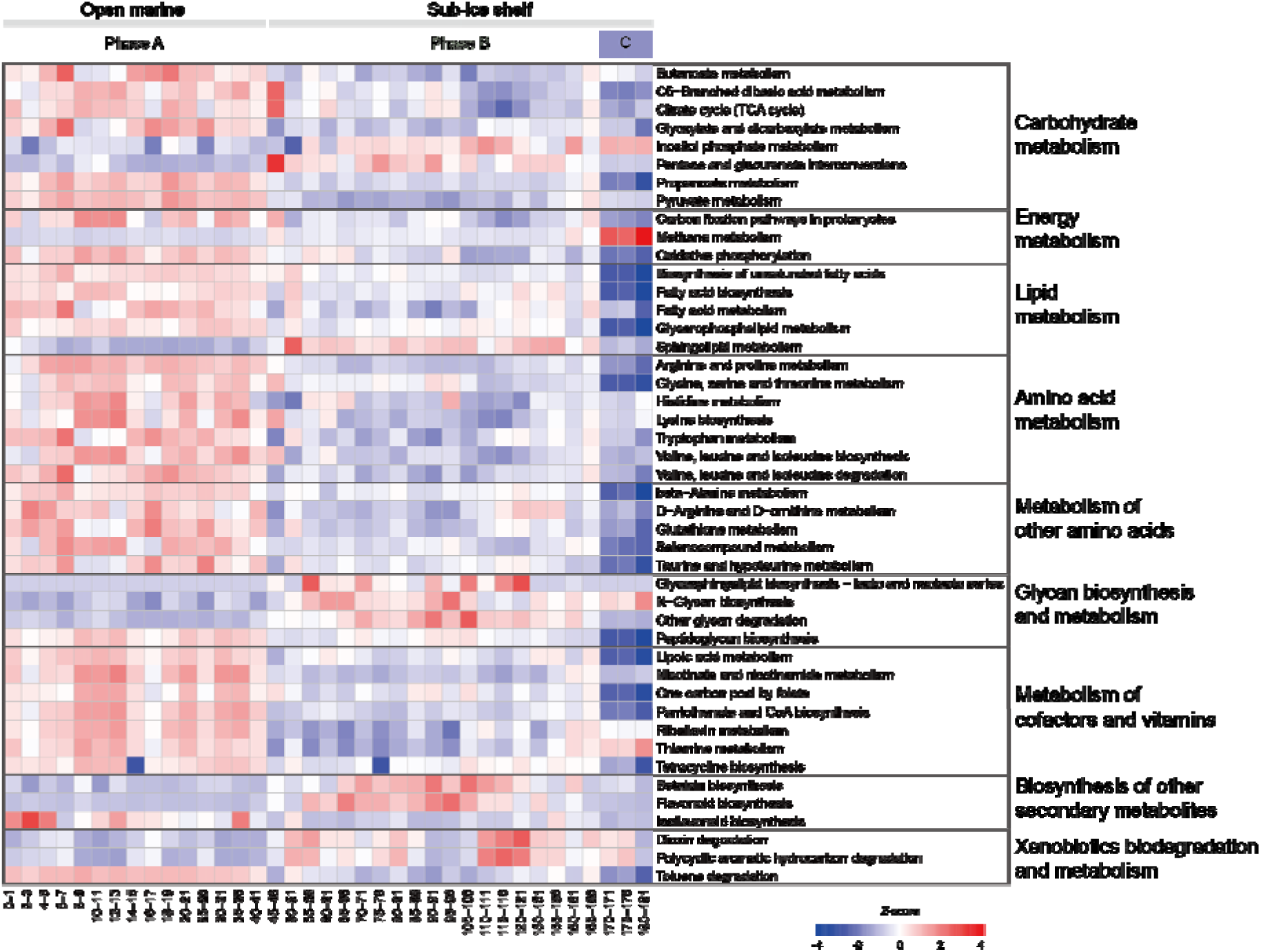
Heatmap of the *Z*-score-normalized relative abundances of the KEGG pathways. Functional profiles of the KEGG pathways were predicted using PICRUSt2 and 45 pathways significantly different between phases A and B‒C are shown.

**Extended Data Fig. 4.**
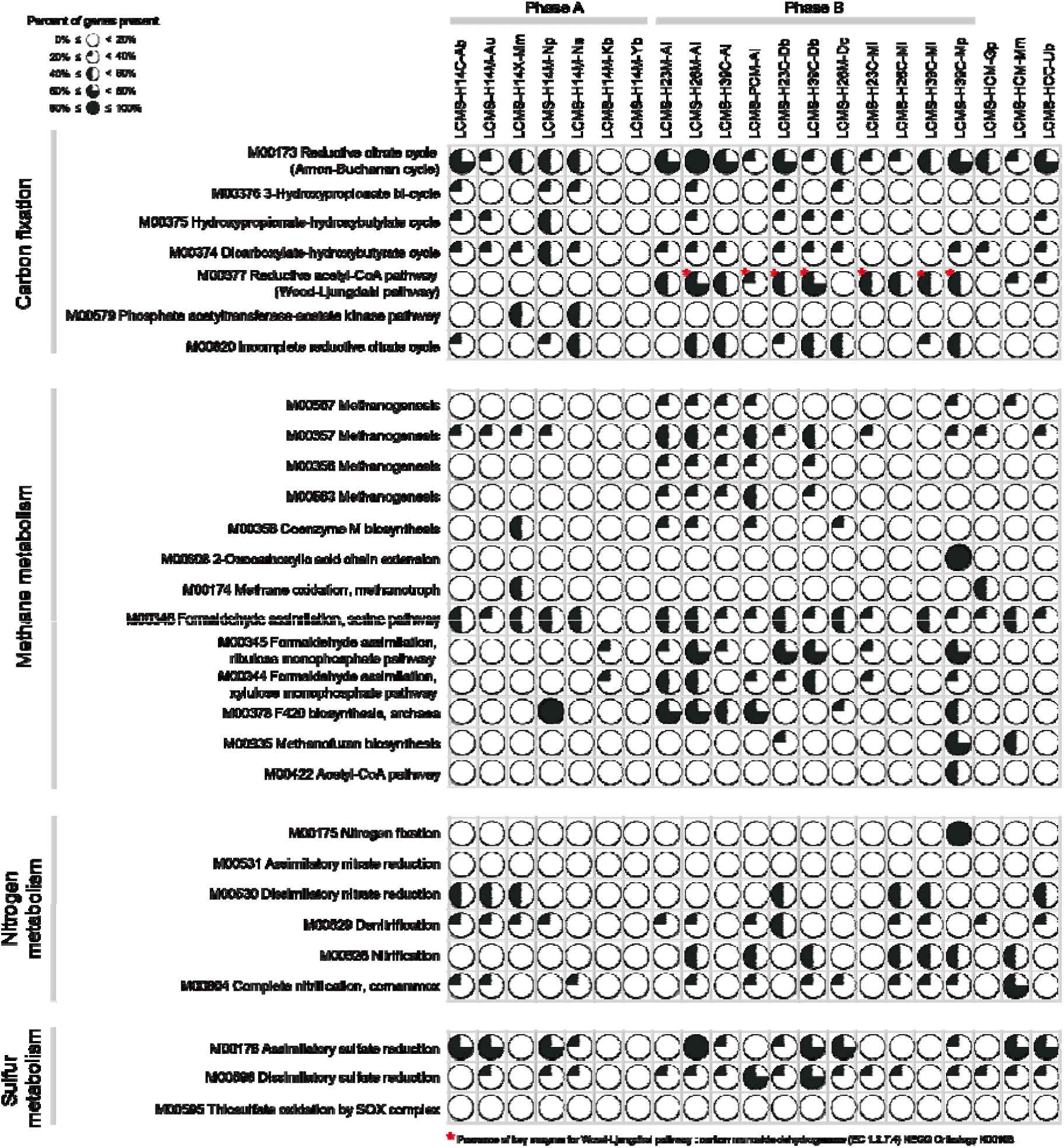
Functional annotation of 21 reconstructed MAGs. Genes involved in carbon fixation, methane metabolism, nitrogen metabolism, and sulfur metabolism were analyzed using BlastKOALA. The percentage of genes present in each KEGG module was calculated using the KEGG mapper. Metagenome-assembled genomes are: LCMS-H14C-Ab (size 1,928,535 bp, N50 2,464 bp, completeness 53.20%, contamination 4.27%, GTDB-tk taxonomic assignment d_Bacteria;p_Acidobacteriota;c_Mor1;o__;f__;g__;s__), LCMS-H14M-Au (1,422,935 bp, 3,382 bp, 56.83%, 2.84%, d_Bacteria;p_Aureabacteria;c__;o__;f__; g__;s__), LCMS-H14X-Mm (2,576,697bp, 2,897bp, 54.23%, 8.21%, d_Bacteria; p_Methylomirabilota;c_Methylomirabilia;o_Methylomirabilales;f__;g__;s__), LCMS-H14M-Ns (1,737,871 bp, 5,868 bp, 79.91%, 3.42%, d_Bacteria;p_Nitrospinota_B;c_2-12-FULL-45-22;o__;f__;g__;s__), LCMS-H14M-Kb (687,815 bp, 10,068 bp, 53.67%, 2.14%, d_Bacteria;p_Patescibacteria;c_Paceibacteria;o_UBA9983;f_UBA2163;g__;s__), LCMS-H14M-Yb (544,689 bp, 8,886 bp, 52.12%, 0%, d_Bacteria;p_Patescibacteria;c_Paceibacteria; o_Paceibacterales;f_UBA10102;g_O2-01-FULL-48-27b;s__), LCMS-H14M-Np (1,440,175 bp, 11,304 bp, 94.66%, 8.314%, d_Archaea;p_Thermoproteota;c_Nitrososphaeria; o_Nitrososphaerales;f_Nitrosopumilaceae;g__;s__), LCMS-H23M-Al (2,665,841 bp, 4,624 bp, 50.55%, 0.91%, d_Bacteria;p_Chloroflexota;c_Anaerolineae;o_Anaerolineales;f_UBA4823_A; g_9FT-COMBO-55-16;s__), LCMS-H26M-Al (3,946,168 bp, 7,416 bp, 83.93%, 4.545% d_Bacteria;p_Chloroflexota;c_Anaerolineae;o_Anaerolineales;f_UBA4823_A;g_9FT-COMBO- 55-16;s__), LCMS-H39C-Al (1,762,169 bp, 2,842 bp, 50.14%, 2.73%, d_Bacteria; p_Chloroflexota;c_Anaerolineae;o_Anaerolineales;f_UBA11858;g__;s__), LCMS-H23C-Db (3,031,859 bp, 1,851 bp, 52.22%, 8.93%, d_Bacteria;p_Desulfobacterota;c_Desulfobacteria; o_Desulfobacterales;f_UBA2156;g_UBA2156;s__), LCMS-H39C-Db (1,976,203 bp, 4,315 bp, 64.64%, 2.10%, d_Bacteria;p_Desulfobacterota;c_BSN033;o_UBA8473;f_UBA8473;g__; s__), LCMS-H23C-Mi (1,292,944 bp, 1,713 bp, 50.92%, 8.59%, d_Bacteria;p_Nitrospirota; c_Thermodesulfovibrionia;o_UBA6902;f_BMS3Bbin08;g__;s__), LCMS-H26C-Mi (1,227,546 bp, 4,839 bp, 58.81%, 7.47%, d_Bacteria;p_Nitrospirota;c_Thermodesulfovibrionia;o_UBA6902; f_BMS3Bbin08;g__;s__), LCMS-H39C-Mi (1,851,896 bp, 3,116 bp, 72.28%, 8.53%, d_Bacteria;p_Nitrospirota;c_Thermodesulfovibrionia;o_UBA6902;f_BMS3Bbin08;g__;s__), LCMS-H26M-Dc (1,610,711 bp, 5,035 bp, 67.49%, 0.99%, d_Bacteria;p_Chloroflexota; c_Dehalococcoidia;o_SM23-28-2;f_RBG-16-64-32;g_RBG-16-64-32;s__), LCMS-H39C-Mp (1,298,657 bp, 3,763 bp, 66.80%, 1.96%, d_Archaea;p_Halobacteriota;c_Methanosarcinia; o_Methanosarcinales;f_Methanoperedenaceae;g_Methanoperedens_A;s__), LCMS-HCC-Ub (1,286,820 bp, 3,489 bp, 63.09%, 2.03%, d_Bacteria;p__;c__;o__;f__;g__;s__), LCMS-HCM-Mm (2,142,260 bp, 8,125 bp, 70.37%, 5,56%, d_Bacteria;p_Methylomirabilota; c_Methylomirabilia;o_Methylomirabilales;f_CSP1-5;g_CSP1-5;s__), LCMS-HCM-Gp (1,933,681 bp, 3,853 bp, 58.08%, 9.74%, d_Bacteria;p_Proteobacteria;c_Gammaproteobacteria;o_BM003; f_BM003;g__;s__), and LCMS-PCM-Al (2,467,625 bp, 3,818 bp, 54.36%, 3.01%, d_Bacteria; p_Chloroflexota;c_Anaerolineae;o_Anaerolineales;f_UBA4823_A;g_9FT-COMBO-55-16;s__).

**Extended Data Fig. 5.**
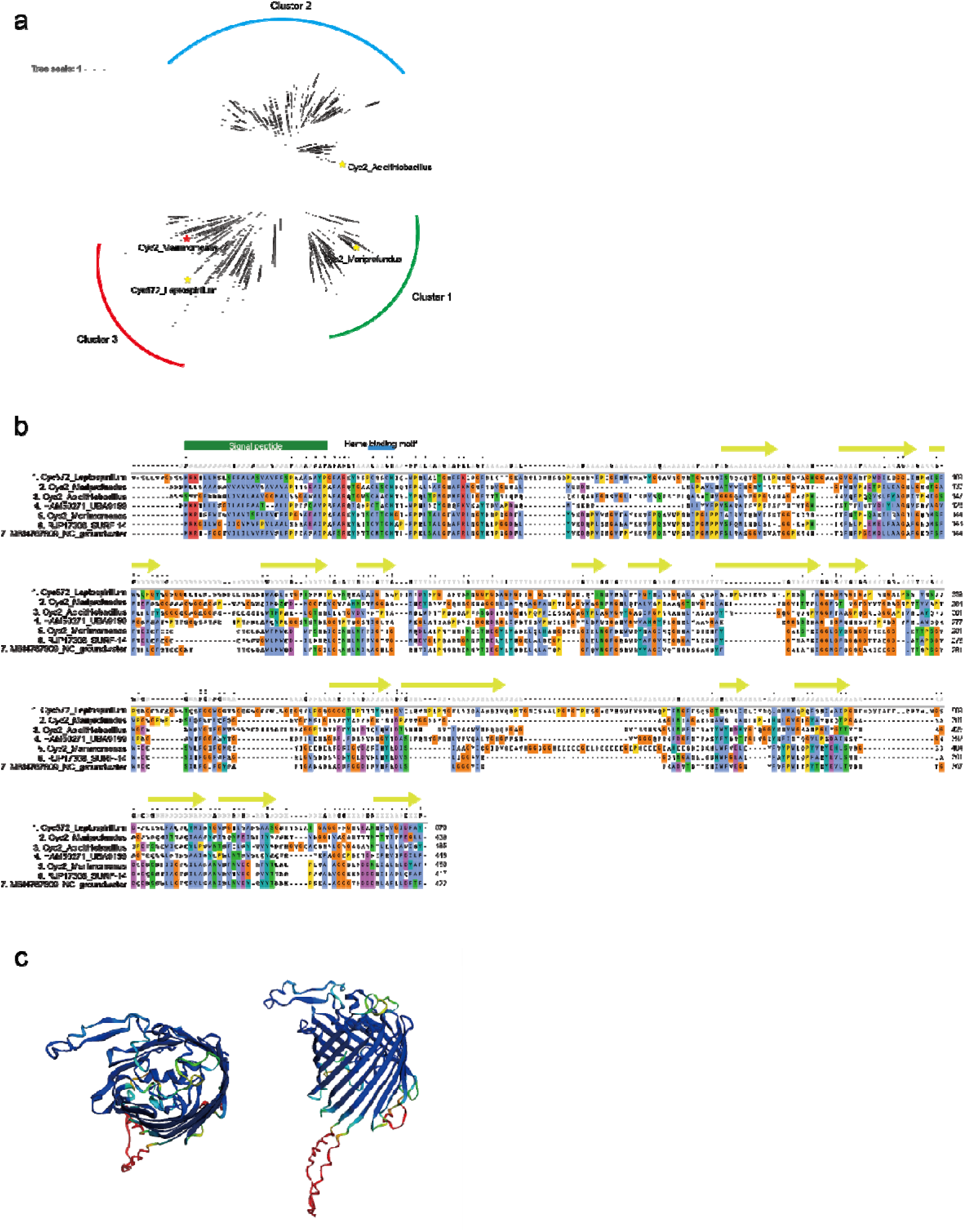
Phylogeny, sequence alignment, and structural prediction of the Cyc2 protein of ‘*Ca*. Mariimomonas ferrooxydans’. **a,** Phylogeny of Cyc2 proteins and their homologs. 1,594 sequences homologous to Cyc2 were retrieved from the database and aligned using ClustalW. A maximum likelihood trees was constructed using RAxML. Yellow stars indicate reference Cyc2 proteins: Cyc2_Acidithiobacillus, *Acidithiobacillus ferrooxidans* ATCC 23270; Cyc572_Leptospirillum, *Leptospirillum* sp. Group II ‘C75’; Cyc2_ Mariprofundus, *Mariprofundus ferrooxydans* PV-1. **b,** Sequence alignment of Cyc2 Fe(II) oxidases and their homologs. The Cyc2 sequence of ‘*Ca*. M. ferrooxydans’ (Cyc2_ Mariimomonas) is aligned along with its closest homologs: RJP17308_SURF_14 from *Deltaproteobacteria* SURF_14 and MBI4767809_NC_groungwater from *Deltaproteobacteria* bacterium (groundwater metagenome). Also shown is HAM50271_UBA9159 from *Nitrospirae* UBA9159. **c,** 3D structure of the Cyc2 protein of ‘*Ca*. M. ferrooxydans’. Two views of the 3D structure predicted using the Alphafold2 algorithm are shown.

**Extended Data Fig. 6.**
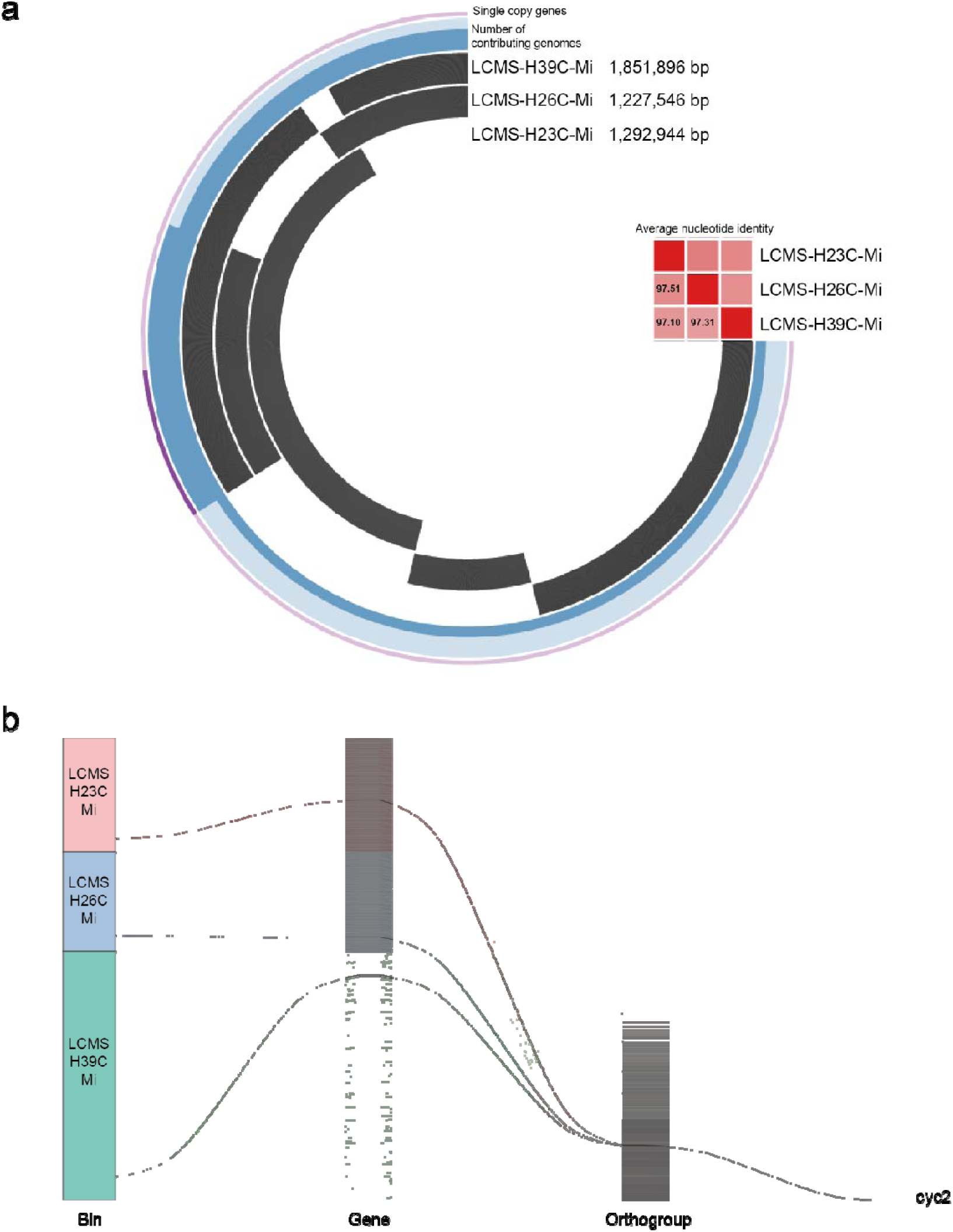
Synteny plots of the three “*Ca*. M. ferrooxydans” MAGs, LCMS-H23C-Mi, LCMS-H26C-Mi, and LCMS-H39C-Mi. **a,** Hierarchical clustering of genes based on their presence in the three MAGs. Single-copy core genes, which are present in all three MAGs, are indicated. Average nucleotide identity (ANI) values between the MAGs are 97.10–97.51%. **b,** Distribution of genes belonging to the three MAGs visualized by an alluvial plot. Genes with consensus sequences were aligned in the same order and grouped into 604 orthogroups, one of which is homologous to *cyc2*. Three genes, which are homologs of *cyc2* and offer phylogenetic inference of these MAGs, stretched out from similar positions in each MAG and converged into the same orthogroup.

**Extended Data Fig. 7.**
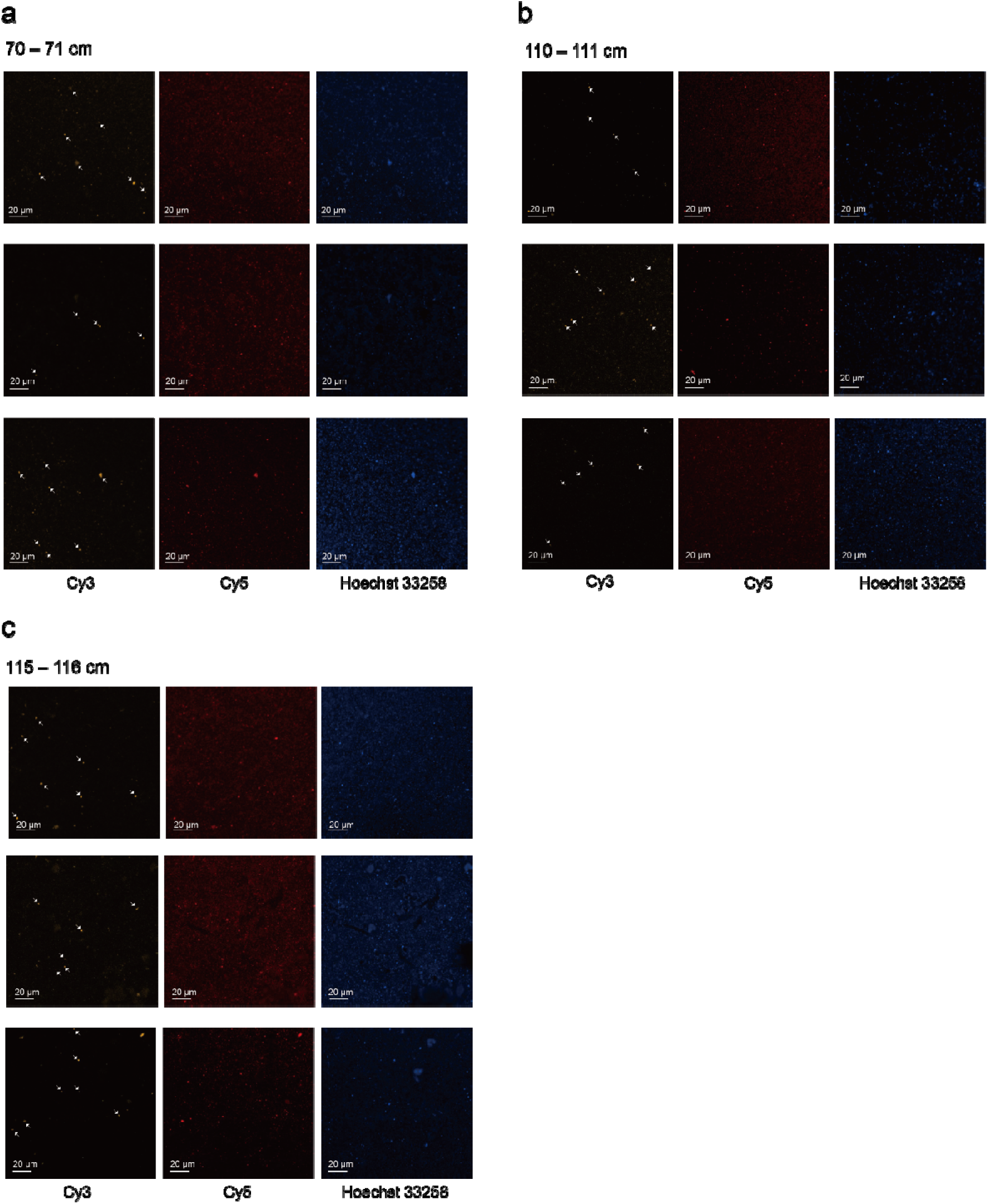
Fluorescence *in situ* hybridization images of the LIS-C sediment samples from 70‒71 cm (a), 110–111 cm (b), and 115–116 cm (c). Oligonucleotide probes target “*Ca*. M. ferrooxydans” (Mari_866F_Cy3, orange) and general bacteria (EUB338_Cy5, red). DNA was stained with Hoechst 33258 (blue). Scale bar, 20 μm.

**Extended Data Fig. 8.**
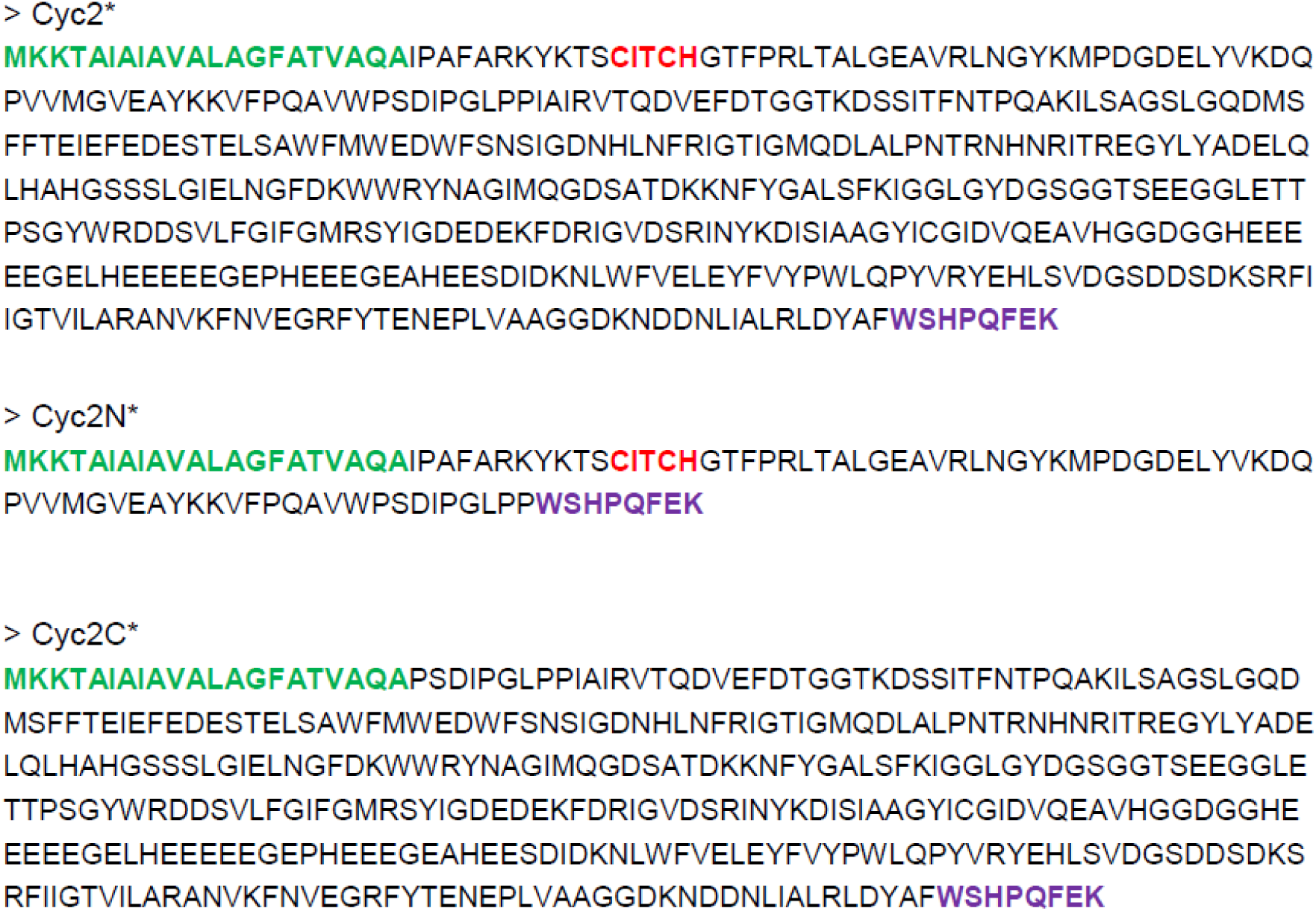
Heterologous expression of ‘*Ca*. M. ferrooxydans’ Cyc2 in *E. coli*. Modified Cyc2 sequences of ‘*Ca*. M. ferrooxydans’ that are used for heterologous expression in *E. coli*. Cyc2*, the full-length Cyc2 sequence with the substituted N-terminal signal peptide of *E. coli* OmpA and the C-terminal Strep-tag II. Amino acid residues shaded in green indicate the signal peptide of *E. coli* OmpA, red the heme-binding motif CXXCH, and purple Strep-tag II. Cyc2N*, the cytochrome c domain of Cyc2 fused to the signal peptide of E. coli OmpA and Strep-tag II. Cyc2C*, the transmembrane β-barrel domain of Cyc2 fused to the signal peptide of E. coli OmpA and Strep-tag II.

