## Supplementary Information for "Who Formed All That Iron?: A Novel Antarctic Chemolithotroph Drives Iron Biomineralization"

**Supplementary Information: Taxonomic Proposal**

**Description of ‘*Candidatus* Mariimomonas’ gen. nov.**

*Mariimomonas* (Ma.ri.imo.mo'nas. L. neut. n. *mare* the sea; L. neut. n. *imum* bottom; Gr. fem. n. *monas* a monad, unit; N.L. fem. n. *Mariimomonas* a monad from sediments in the seabed).

The DNA G+C content of the only representative type species is 41.2% according to the LCMS-H39C-Mi sequence. Based on the 16S rRNA gene sequence and phylogenomic analyses, the provisional genus belongs to the family ‘*Candidatus* Mariimomonadaceae’, the order ‘*Candidatus* Mariimomonadales’, the class *Thermodesulfovibrionia* Umezawa *et al*. 2021^1^, and is clearly different (≤87% 16S rDNA identity) from all other genera with validly published names and *Canditatus* taxa comprising *Thermodesulfovibrionia*. There are no reference genomes closely related to this genus, and genomes of the type species share just 72% average nucleotide identity to BMS3Bbin08 (*Nitrospirae* Clade B) of an as-yet-uncultivated chemosynthetic colonizer in seafloor massive sulphide deposits^2^. Obligately anaerobic and chemolithoautotrophic.

The type species is ‘*Candidatus* Mariimomonas ferrooxydans’.

**Description of ‘*Candidatus* Mariimomonas ferrooxydans’ sp. nov.**

*Mariimomonas ferrooxydans* (ferr.ox'y.dans. L. n. *ferrum* iron; Gr. adj. *oxys* sour; N.L. v. *oxydo* to sour, oxidize; N.L. part. adj. *ferrooxydans* iron-oxidizing).

The species occupies an important place, with markedly high centralities, in the co-occurrence sub-network of phase B–C (layers 45‒181 cm) in a whole round core sample of the marine sediment (EAP13-GC16B; 236 cm below the seafloor at 324 m below the sea level in the approximately 8.7-km-wide basin), mostly consisting of laminated facies deposited during the Holocene, beneath the Larsen C Ice Shelf on the northern part of the embayment (66° 3.89832′ S, 60° 27.69212′ W) in Antarctica. Cells in the sediment samples were visualized via fluorescence *in situ* hybridization using a 16S rRNA-targeted and fluorophore-labelled probe. Based on the draft metagenome-assembled genomes (MAGs), the provisional species was predicted to encode the pathways for dissimilatory sulphate reduction and nitrate reduction to ammonia. It is able to utilize acetate and formate as electron donors. Harbouring a Wood–Ljungdahl pathway that involves carbon monoxide dehydrogenase, the species possesses the metabolic potential to oxidize ferrous iron through the Cluster 3-type Fe(II) oxidase Cyc2.

The designated type materials are the 16S rRNA gene of OTU7 and the MAG LCMS-H39C-Mi, which were deposited in GenBank under BioProject PRJNA792354. Based on medium-quality LCMS-H39C-Mi, its G+C content is 41.2% and estimated genome size is >1.85 Mb. The GenBank/EMBL/DDBJ accession numbers of the partial 16S rRNA gene sequence and the draft genome sequences of LCMS-H23C-Mi, LCMS-H26C-Mi, and LCMS-H39C-Mi (97.10–97.51% nucleotide identities) are ON025791, SAMN24803906, SAMN24803907, and SAMN24803905, respectively.

**Description of ‘*Candidatus* Mariimomonadaceae’ fam. nov.**

*Mariimomonadaceae* (Ma.ri.imo.mo.na.da.ce'ae. N.L. fem. n. *Mariimomonas* type genus of the family; L. suff. -*aceae* ending to denote a family; N.L. fem. pl. n. *Mariimomonadaceae* the family of the genus *Mariimomonas*).

The family ‘*Candidatus* Mariimomonadaceae’ is within the order ‘*Candidatus* Mariimomonadales’ and encompasses Gram-negative bacteria whose DNA was retrieved from marine sediments. The family was established on the basis of 16S rRNA gene- and genome-based phylogeny, and is a member of the order ‘*Candidatus* Mariimomonadales’ in the class *Thermodesulfovibrionia*. Currently, the family comprises the genus ‘*Candidatus* Mariimomonas’. The description of the family is the same as described for the genus ‘*Candidatus* Mariimomonas’. The type genus is ‘*Candidatus* Mariimomonas’.

**Description of ‘*Candidatus* Mariimomonadales’ ord. nov.**

*Mariimomonadales* (Ma.ri.imo.mo.na.da.a'les. N.L. fem. n. *Mariimomonas* type genus of the family; L. suff. -*ales* ending to denote an order; N.L. fem. pl. n. *Mariimomonadales* the order of the genus *Mariimomonas*).

The order ‘*Candidatus* Mariimomonadales’ is within the class *Thermodesulfovibrionia* and encompasses Gram-negative bacteria whose DNA was retrieved from subsurface environments. The family was established on the basis of 16S rRNA gene- and genome-based phylogeny, and is a member of the class *Thermodesulfovibrionia* in the phylum *Nitrospirae* Garrity and Holt 2001^3^ (*Nitrospirota*^4^ ). According to the MAG sequences of LCMS-H23C-Mi, LCMS-H26C-Mi, LCMS-H39C-Mi, SZUA-198 (GCA_003229575.1), BMS3Bbin08 (GCA_002897775.1), DSMQ01 (GCA_011047655.1), JDFR-81 (GCA_002011735.1), SURF-23 (GCA_003599425.1), SURF-11 (GCA_003599505.1), SURF-45 (GCA_003599275.1), JAADFS01 (GCA_013152795.1), UBA6902 (GCA_002451135.1), JACNKH01 (GCA_014382155.1), BMSAbin06 (GCA_002897895.1), BMSAbin09 (GCA_002897915.1), and Glo-13 (GCA_003354025.1), members of the order commonly encode the Wood–Ljungdahl pathway, a nitrate reductase, and the Dsr–Apr–Sat system. Their gene contents suggest that the *Candidatus* order maintains a chemolithoautotrophic lifestyle. Currently, the order comprises the family ‘*Candidatus* Mariimomonadaceae’. The description of the order is the same as described for the family ‘*Candidatus* Mariimomonadaceae’. The type genus is ‘*Candidatus* Mariimomonas’.
